# Aberrant 5-Methylcytosine tRNA Modification Disrupts Proteostasis and Exacerbates Age-Related Osteoporosis

**DOI:** 10.1101/2025.11.04.686482

**Authors:** Zhuolin Peng, Ning Peng, Mengzhu Yuan, Feng Yan, Tiejian Jiang, Mingsheng Ye, Ye Xiao

## Abstract

Age-related protein aggregation drives senile osteoporosis. Aberrant tRNA modifications exacerbate the progression, yet mechanisms linking these to bone loss remain unclear. In this study, we identify *Nsun2* (m^5^C methyltransferase) as key regulator: age-dependent *Nsun2* downregulation in BMSCs reduces m⁵C levels of tRNAs, destabilizing tRNAs and impairing translation efficiency of specific transcripts. This directly disrupts the protein synthesis of molecular chaperone and pro-osteogenic factors, accelerating misfolded protein aggregates and activating unfolded protein response, inducing BMSCs senescence and impairing osteogenesis. Mice specifically depleted of *Nsun2* exhibited reduced bone mass, whereas mice overexpressing *Nsun2* alleviated age-associated bone loss. Notably, the exacerbated protein aggregation and bone mass loss in *Nsun2*-deficient mice were ameliorated following treatment with the molecular chaperone activator-ML346. Remarkably, ML346 administration proved sufficient to reverse age-related functional deficits in aged mice. Overall, our findings demonstrate that aberrant tRNA-m^5^C modification alters protein synthesis and induces proteostasis collapse, which constitute a novel contributor to the pathogenesis of senile osteoporosis. Additionally, reduction of protein aggregation through the activation of molecular chaperones presents a promising therapeutic strategy for this disease.

## Introduction

Prevailing theories about the mechanisms of age-related diseases identify the gradual disruption of cellular protein homeostasis (proteostasis) as a central factor, a process characterized by the accumulation of protein aggregates(1, 2). Osteoporosis is defined as an age-associated skeletal disorder resulting from disrupted bone homeostasis, marked by an imbalance between bone formation and resorption(3). Recent researchs have reported that disruption of proteostasis during aging leads to an imbalance in bone homeostasis, which is a major factor in senile osteoporosis(4). However, the molecular mechanisms driving proteostasis disruption and the contribution of protein aggregates to the pathogenesis of senile osteoporosis is not fully understood.

Proteostasis refers to a dynamic homeostatic state in which protein synthesis, folding, and degradation are precisely regulated to maintain proteome integrity and ensure normal cellular function(5, 6). The maintenance of proteostasis depends on the proteostasis network (PN), which consists of molecular chaperones and two major protein degradation systems: the ubiquitin-proteasome system (UPS) and autophagy (7).In healthy cells, molecular chaperones facilitate correct protein folding by binding and stabilizing unstable conformations, while also recognizing and interacting with misfolded proteins to prevent aberrant aggregation and promote refolding or disaggregation(8). If misfolded proteins escape chaperone-mediated surveillance, they are typically targeted for degradation via the UPS or autophagy pathways(9). This quality control mechanism is critical for preserving cellular function and preventing protein aggregation. During aging, dysfunction of molecular chaperones and protein degradation systems leads to protein aggregation, which subsequently induces certain age-associated degenerative diseases(8, 10, 11). In osteoporosis progression, UPS dysfunction and age-related increase in osteoclast autophagy promote osteoclastogene-sis through mediation of RANKL-induced NF-κB signaling activation. Conversely, osteocyte/osteoblast autophagy deficiency by inducing the unfolded protein response (UPR) and sustaining endoplasmic reticulum (ER) stress, along with UPS-mediated ubiquitin degradation of pro-osteogenic proteins, collectively impair age-related bone formation(4, 12–14). Thus, dysregulation of protein degradation may be regarded as a mechanistic explanation for how ageing disrupts proteostasis and drives age-related osteoporosis. Notably, while the role of protein degradation in osteoporosis progression has been extensively characterized, the regulatory mechanisms connecting age-related alters in protein synthesis to proteostasis imbalance and their specific impact on bone loss remain unclear.

Several studies have reported a significant reduction in global protein synthesis rates across various tissues in aging mice and rats(15). This decline may reflect an adaptive response to aging that is potentially beneficial for longevity; however, it may also reduce the production of key functional proteins, molecular chaperone proteins(16). Previous studies have demonstrated that molecular chaperone synthesis is essential for maintaining proteostasis(2, 17). Specifically, the molecular chaperone protein declines, which may decrease protein folding capacity and increase the accumulation of unfolded or misfolded proteins in aged cells(18–20). This phenomenon may be explained by the observation that aging weakens the correlation between transcript levels and their corresponding protein products. Notably, as core elements of the translation machinery, transfer RNAs (tRNAs) undergo precise aminoacylation to form aminoacyl-tRNAs-critical intermediates that decode mRNA codons during ribosome-mediated protein synthesis. Aging is commonly accompanied by an increase in tRNA-derived fragments (tRFs) and a decrease in the abundance of specific tRNAs(21). tRNA modification,including methylation, hydroxylation, acetylation, and deamination, serve to protect tRNAs from cleavage into tRFs, thereby maintaining tRNA abundance and ensuring efficient translation(22, 23). These modifications are critical for enhancing the accuracy of codon recognition, improving decoding efficiency, and maintaining the overall efficacy of translation. Loss of tRNA modifications compromises their structural stability, resulting in ribosomal stalling at specific codons and dysregulated protein expression(24, 25). Furthermore, it impairs translational fidelity, which can trigger mRNA degradation and promote protein misfolding, which has been closely associated with various diseases including type2 diabetes, neurodegenerative disorders and tumorigenesis(26). Among these epigenetic modifications, methylation is the most prevalent post-translational modification. Notably, the 5-methylcytosine (m⁵C) modification, in particular, serves as a critical molecular bridge that directly couples protein synthesis machinery to cellular metabolic pathways(27). This modification is catalyzed by at least eight highly conserved enzymes, including NSUN1-7 and DNMT2. Among these, NSUN2 has been the most relevant methyltransferase(28). Utilizing S-adenosylmethionine (SAM) as the methyl donor, NSUN2 specifically methylates select coding and non-coding RNAs, as well as the anticodon loops and variable regions of tRNAs, thereby protecting tRNAs from cleavage(29). Notably, previous studies have shown that external stress can suppress *Nsun2* expression, resulting in reduced tRNA m⁵C methylation, decreased tRNA stability and ultimately impaired protein synthesis and stem cell function(30–32). This evidence indicates that *Nsun2*-mediated tRNA m⁵C modification is tightly coupled to protein synthesis. It is well-established that the imbalance in bone homeostasis is a major factor in senile osteoporosis. However, whether *Nsun2*-mediated tRNA m^5^C modification regulates bone homeostasis by influencing protein synthesis and proteostasis remains unclear.

In this study, we reveal that age-associated cellular stress suppresses *Nsun2* expression, reduces tRNA m^5^C methylation, and impairs molecular chaperone activity or pro-osteogenic protein synthesis. This cascade of molecular dysfunctions disrupts proteostasis, leading to protein misfolding/aggregation, activation of UPR, and sustained ER stress - thereby induces age-associated osteoporosis. Conversely, overexpression of *Nsun2* restores m^5^C methylation, restore efficient protein synthesis, and rescue age-related protein aggregation, and mitigate differentiation imbalance.

These findings elucidate the functional significance of *Nsun2*-dependent m^5^C tRNA modification in bone homeostasis, uncover the underlying molecular and cellular mechanisms, and highlight its potential as a therapeutic target for osteoporosis.

## Results

### Age-dependent dysregulation of tRNA m^5^C methylation and translational efficiency in senescent BMSCs

Transfer RNAs (tRNAs) are a fundamental component of the translation machinery and serve as adapter molecules for coordinating mRNA translation and protein synthesis. To date, more than 170 tRNA modifications have been identified, which are crucial for maintaining tRNA stability, activity, and ensuring accurate and efficient protein synthesis(33, 34) (Fig. 1A). Dysregulation of tRNA modification is implicated in several distinct age-associated diseases, including neurological disorders, mitochondrial disease, and cancer(35). Senile osteoporosis is defined as an age-related bone metabolic disorder. However, it remains unclear whether aberrant tRNA modification is involved in the progression of the disease. To investigate the hypothesis, we first characterized the tRNA modification landscape via LC-MS/MS in bone marrow mesenchymal stromal cells (BMSCs) isolated from 24-month-old mice (a good model for senile osteoporosis). BMSCs, being multipotent stem cells, exhibit age-related functional decline, which serves as the primary driver for senile osteoporosis (36). We found that several reported tRNA modifications associated with translational accuracy and tRNA stability-including 5-methylcytosine (m^5^C), N1-methyladenosine (m^1^A), N6-methyladenosine (m^6^A), N7-methylguanosine (m^7^G), N1-methylguanosine(m^1^G) and N6-threonylcarbamoyladenosine (t^6^A)-were identified (Fig. 1A and 1B). Among these, 5-methylcytosine (m^5^C) emerged as the most prevalent modification (Fig. 1B). In tRNA, m^5^C is one of the most abundant tRNA modifications and is often required for normal development through promoting tRNA stability and efficient protein synthesis. Aberrant m^5^C hypermethylation alters stem cell functions and causes neurodevelopmental and mitochondrial disorders(30, 31, 37). To identify age-dependent changes in m^5^C levels, the isolated BMSCs from 2-month-old (young) and 24-month-old mice (old) were cultured and performed subjected to bisulfite sequencing (Bis-seq). This sequencing results revealed that cytosine-5 methylation levels at the junction between the variable loop (VL) and the TΨC arm (positions 48/49, Ala-CGC, Gln-TGG, Gly-GCC, Lys-CTT, Thr-CGT, Thr-AGT, Val-CAC and Val-TAC), were significantly decreased in old BMSCs compared to young BMSCs (Fig. 1C-D). These changes were validated by Northern blotting (Fig. 1E), which suggests that aberrant m^5^C hypermethylation of tRNA may be tightly associated with BMSC senescence. Previous studies indicated that reducing tRNA m^5^C modification destabilizes tRNAs, leading to decreased abundance(37). Indeed, we found that the abundance of m^5^C tRNAs was downregulated in old BMSCs compared to young BMSCs, suggesting that the stability of hypo-modified tRNAs was lost in senescent BMSCs (Fig. 1F). Besides, we also found that the abundance of some non-m^5^C-modified tRNAs was decreased, suggesting that other RNA-modifying enzymes, beyond those involved in m^5^C modification, may regulate tRNA stability or abundance in senescent cells. Given the positive correlation between cytosine-5 RNA methylation and protein synthesis(31, 32), we hypothesize that protein synthesis rate declines in senescent BMSCs. To examine this, we isolated and cultured senescent and young BMSCs with azide-modified molecules, after which the modified proteins were extracted and pulled down using Alkyne PEG Biotin via a Click-iT Reaction. Western blot analysis revealed a significant reduction in nascent polypeptides (NP) in senescent BMSCs compared to young cells, which consistent with the age-dependent decline in protein synthesis rates during aging (Fig. 1G)(38, 39). As the mRNA codon decoder during protein synthesis, alterations in tRNA abundance or their chemical/structural modifications will directly impact ribosomal polypeptide elongation rates and translation efficiency, consequently inducing translational pauses that may have wide implications for protein homeostasis, protein quality control, and disease(24). We next investigated whether alterations in the abundance of m⁵C-modified tRNAs influence translational efficiency and ribosomes stalling at codons decoded by tRNAs. To examine this, we conducted genome-wide ribosomal footprinting assays (Ribo-seq) (Fig. 1H), the trinucleotide periodicity of ribosomes and codon occupancy were estimated using revised riboWaltz4 package and translational efficiencies were determined as the ratio of (normalized abundance determined by ribosome profiling)/(normalized abundance determined by RNA-seq) as previously repored(40, 41). Expectedly, senescent BMSCs exhibited a significant increase in both dwell time (DT) and E-site/P-site (the site holding the tRNA associated with the growing polypeptide chain) occupancy compared to young cells (Fig. 1I and 1J). Translational efficiency (TE) analysis revealed that 806 mRNAs exhibited a great increase in TE, while 848 mRNAs showed a decrease in TE, with similar counts of mRNAs displaying either increased or decreased TE (Fig. 1K). Gene ontology analysis revealed that mRNAs with reduced TE in senescent BMSCs-functioning in key biological processes including cellular calcium ion homeostasis, DNA repair, skeletal system development, Notch signaling pathway and stem cell differentiation-were significantly enriched (Fig. 1L). Increased codon occupancy may lead to translational pausing. As we expected, analysis of codon pausing using PausePred software revealed that codon pausing was significantly elevated in genes associated with skeletal system development and BMSCs differentiation (*Col1a1*, *Runx2*, *Lrp1*, *Spp1*, *Yap1*, *Foxp1,* etc) in old BMSCs compared to young cells (Fig. 1M). This pattern suggests that aberrant 5-Methylcytosine tRNA modification alters tRNA abundance and translational efficiency, which may selectively regulate the expression of pro-osteogenic proteins, thereby influencing the function of BMSCs and cellular senescence.

**Figure 1.**
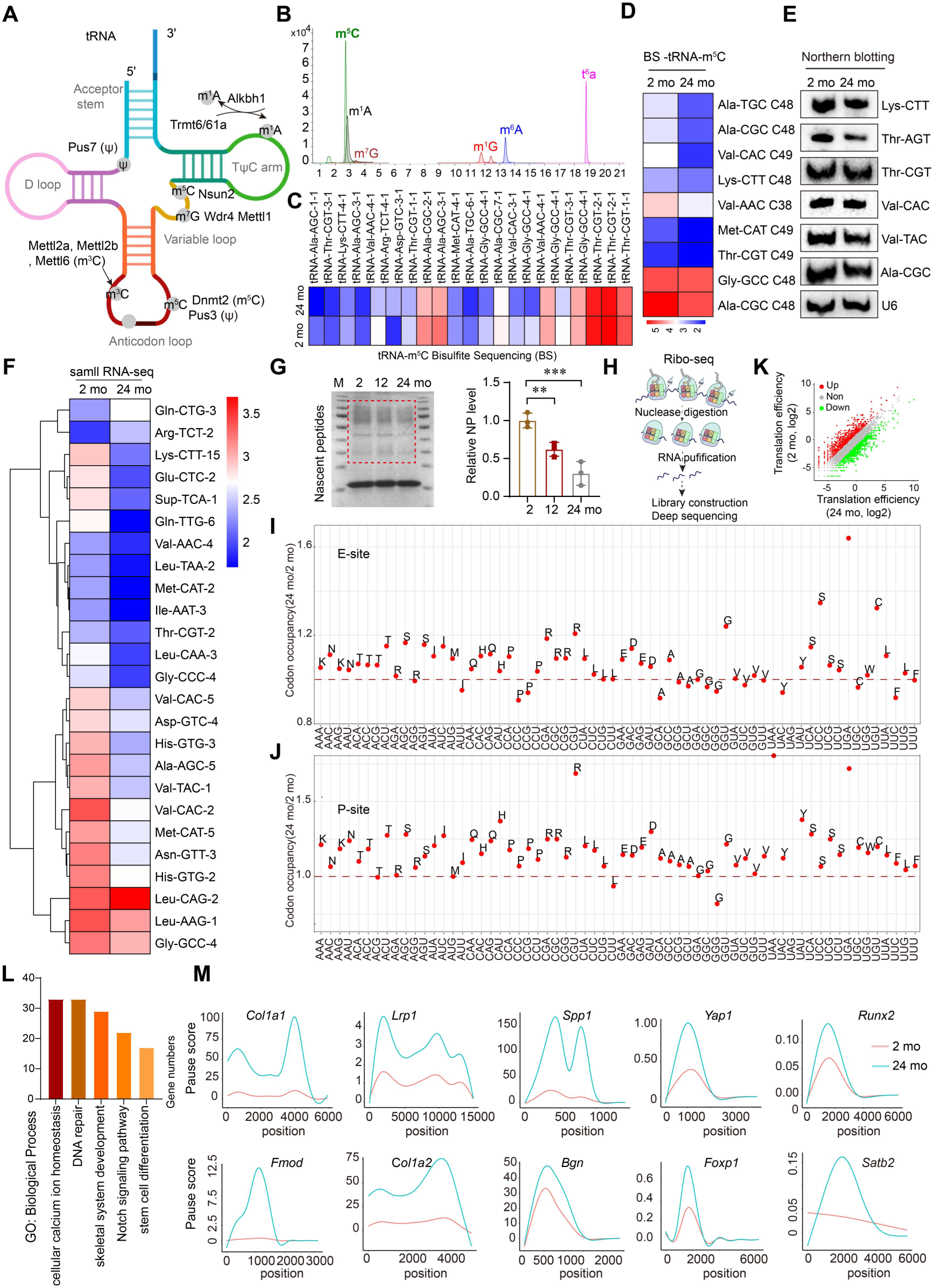
Age-dependent dysregulation of tRNA m^5^C methylation and translational efficiency in senescent BMSCs. (A) Schematic diagram of tRNA secondary structure and modification types. (B) Total ion chromatogram of LC-MS/MS analysis of modified tRNA ribonucleosides. (C) Heatmap of m^5^C-modified tRNAs in BMSCs from 2-, and 24-month-old mice. The colour indicates the relative intensity of methylation changes. (D) Heatmap of m^5^C-methylated cytosines at the VL junction (C48, C49) in candidate tRNAs from BMSCs of 2-month-old and 24-month-old mice. (E) Representative northern blotting analysis of candidate tRNA levels in 2-, and 24-month-old mice, and U6 snRNA was used as a loading control. (F) Heatmap of tRNAs abundance in the BMSCs from 2-, and 24-month-old mice, with each colour shows the relative intensity of changes in tRNA abundance. (G) Representative Western blot images of total nascent peptides and quantification of relative nascent peptides level of BMSCs from 2 -, 12 -, and 24-month-old mice. (H) Schematic diagram illustrating the principle of ribosomal footprinting assays (Ribo-seq). (I) Ribosome occupancy at individual codons at E site. Plots represent the relative ribosome protected fragment signals from 24-month-old BMSCs relative to 2-month-old BMSCs. (J) Ribosome occupancy at individual codons at P site. (K) Scatterplot of translation efficiency (TE) in BMSCs from 24-month-old mice versus 2-month-old mice. (L) Gene ontology analysis revealed enrichment of biological processes among the TE downregulated genes of BMSCs from 24-month-old mice compared to 2-month-old mice. (M) Relative ribosome pausing scores of osteogenic differentiation related genes (*Col1a1*, *Lrp1*, *Spp1*, *Yap1*, *Runx2*, *Fmod*, *Col1a2*, *Bgn*, *Foxp1*, *Satb2*) between 2- and 24-month-old mice. Data shown as mean ± SEM. \*\**P* < 0.01, \*\*\**P* < 0.001 by and one-way ANOVA (G).

### *Nsun2* depletion disrupts tRNA stability and global translation via reduced m^5^C methylation

Cytosine-5 (m^5^C) tRNA methylation has been characterized across diverse tRNA species at multiple positions, with its formation is mediated by at least eight highly conserved enzymes-including NSUN1-7 and DNMT2(42). Among these, three currently identified cytosine-5 tRNA methyltransferases in higher eukaryotes are NSUN2 and DNMT2. RNA-sequencing analysis of BMSCs from young and old mice revealed a statistically significant downregulation of *Nsun2* in aged BMSCs compared to young cells (Fig. 2A). This downregulation was further substantiated by RT-qPCR, Western blot, and immunofluorescence analyses, which specifically confirmed the reduction in *Nsun2*. While the expression of *Dnmt2* remained unchanged in senescent BMSCs. (Fig. 2B-D). Leptin receptor-positive (LepR^+^) cells in the bone marrow constitute a major subset of multipotent mesenchymal stromal cells(43). To measure NSUN2 expression in *vivo*, we collected femur samples from 2-month-old and 24-month-old mice and performed co-staining with LepR and Nsun2 antibodies. Immunofluorescence staining of femoral sections demonstrated that Nsun2 was predominantly expressed in LepR⁺ cells and markedly reduced in aged mice (Fig. 2E). These findings indicate that age-dependent downregulation of *Nsun2* may serve as a mechanistic basis for the age-associated loss of tRNA m^5^C methylation. To directly investigate the regulatory role of *Nsun2* in tRNA m^5^C methylation, we knocked down *Nsun2* expression in primary BMSCs using a small interfering RNA (siRNA) interference system. RT-qPCR analysis revealed that the mRNA level of *Nsun2* is significantly reduced in the si*Nsun2*-treated group compared with the negative control (NC) group (Fig. 2N). This reduction suggests successful knockout of *Nsun2* in BMSCs. The liquid chromatography-tandem mass spectrometry (LC-MS/MS) was employed to quantify total 5-methylcytidine levels in purified tRNA samples from si*Nsun2*-treated and NC group. The analysis revealed a global reduction in cytosine-5 methylation (fold change = 2.964) in the si*Nsun2*-treated group relative to the NC group (Fig. 2F). To comprehensively understand the site-specific cytosine-5 tRNA methylation pattern, RNA bisulfite sequencing was performed. The sequencing analysis revealed that cytosine-5 methylation levels at C38 in the anticodon loop (tRNA Val and Asp), and at the junction between the variable loop (VL) and the TΨC arm (positions 48/49/50, Thr, Met, Arg, Val, Pro, Phe, Ala, Glu, Gly, Asp and Lys), were significantly decreased in *Nsun2*-deficient cells compared to control cells (Fig. 2G). Given that loss of *Nsun2* markedly reduces m⁵C modification and disrupts tRNA stability(37, 44), we next asked whether the hypo-modified tRNAs would decrease the abundance of certain tRNAs. Using small RNA-seq, we identified that a total of 32 certain tRNAs-most of which were m⁵C-modified and exhibited a fold change < 0.75-were decreased in the si*Nsun2*-treated group relative to the NC group (Fig. 2H). This is consistent with a previous study reporting that loss of m^5^C tRNA modification affects tRNA stability(37). Functioning as an amino acid transporter, we speculate that alterations in tRNAs abundance may signal changes in bulk amino acid content. To examine this, we used an unbiased metabolomic approach to profile 19 of the 20 canonical amino acids, which is consistent with a previous study(37). Obviously, there was a significant increase in lysine, glycine and tyrosine levels in the si*Nsun2*-treated group relative to the NC group (Fig. 2I). In contrast, the remaining 16 amino acid levels showed minimal differences between the si*Nsun2*-treated and NC groups. Combining three independent sequencing assays, we observed that *Nsun2* depletion leads to more pronounced abundance deficits across multiple tRNA-Lys and tRNA-Gly isodecoders, which were further confirmed by northern blot analysis (Fig. 2 J-K). A previous study reported that *Nsun2* deficiency results in significant deficits in tRNA-Gly^GCC^ and tRNA-Gly^CCC^ isodecoders in the cortex. Our data further support this finding by isoacceptor-specific RT-qPCR, indicating that *Nsun2* deficiency indeed disrupts tRNA^-Gly^ and tRNA^-Lys^ abundance and may reduce expression of Glycine-rich and Lysine-rich proteins (Fig. 2 L).

**Figure 2.**
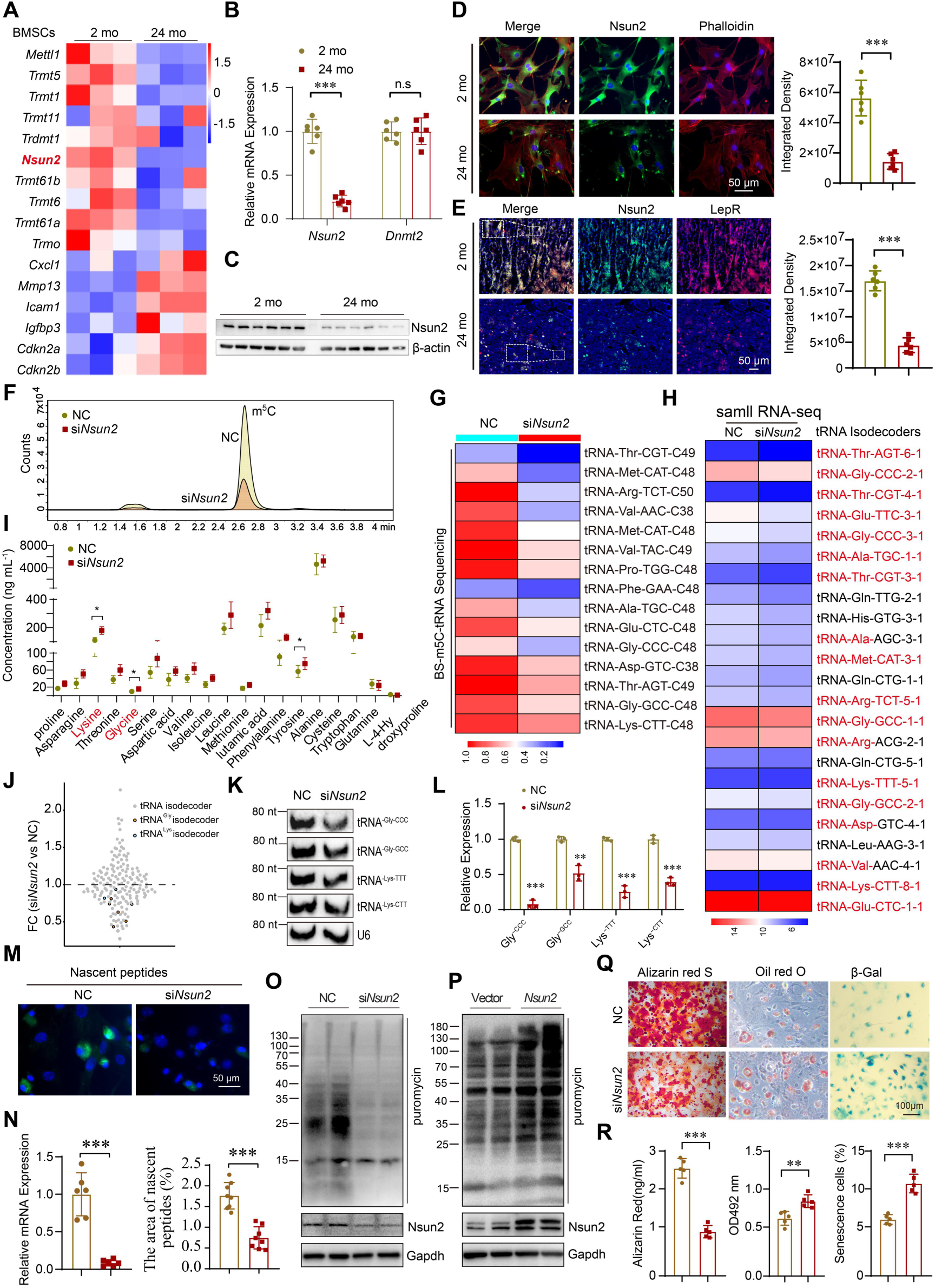
*Nsun2* depletion disrupts tRNA stability and global translation via reduced m^5^C methylation. (A) Heatmap of RNA-seq analysis showing expression changes of methylation-modifying enzymes in BMSCs from 2- and 24-month-old mice. (B) RT-qPCR analysis of *Nsun2* and *Dnmt2* expression levels in BMSCs from 2- and 24-month-old mice. n=6. (C) Representative Western blot images of NSUN2 expression in BMSCs from 2- and 24-month-old mice. n=6. (D) Representative immunofluorescence images and quantification of Nsun2 (green) and Phalloidin (red) staining in femurs of 2- and 24-month-old mice. Scale bar: 50 μm n=6. (E) Representative immunofluorescence images and quantification of NSUN2 (green) and LEPR (red) staining in femurs of 2- and 24-month-old mice. Scale bar: 50 μm n=6. (F) Ridge plot of methylation ratio density in primary BMSCs transfected with negative control RNA or siRNA of *Nsun2*. (G) Heatmap of m^5^C-methylated cytosines at the anticodon loop (C38) and VL junction (C48, C49) in candidate tRNAs from BMSCs of the *siNsun2*-treated group versus the NC group, generated via Bisulfite sequencing (Bis-seq). (H) Small RNA sequencing of tRNA abundance in si*Nsun2*-treated group and the NC group. (I) Gas Chromatography-Tandem Mass Spectrometry (GC-MS/MS) analysis of free amino acid concentrations in si*Nsun2*-treated group and the NC group. (J) tRNA sequencing of primary BMSCs in si*Nsun2*-treated group and the NC group. Note decreased expression in 5 tRNA^Gly^ isodecoders (yellow) and 3 tRNA^Lys^ isodecoders (blue) among total isodecoders (gray). (K) Representative northern blotting analysis of candidate tRNA expression levels in si*Nsun2*-treated group and the NC group, and U6 snRNA was used as a loading control. (L) RT-qPCR analysis of tRNA^Gly-GCC^, tRNA^Gly-CCC^, tRNA^Lys-CTT^ and tRNA^Lys-TTT^ isodecoders expression levels in si*Nsun2*-treated group and the NC group. n=3. (M) Representative immunofluorescence images of nascent proteins (green) and DAPI (blue) staining in si*Nsun2*-treated group and the NC group incubated with OP-puromycin. (N) RT-qPCR analysis (left) of *Nsun2* expression levels and quantification (right) of the percentage of NP area in si*Nsun2*-treated group and the NC group. n=6. (O) si*Nsun2*-treated group and the NC group were incorporated with puromycin (10 μg/ml) for 20 min and subjected to immunoblot analysis. (P) BMSCs transfected with vector or *Nsun2* plasmid were incorporated with puromycin (10 μg/ml) for 20 min and subjected to immunoblot analysis. (Q) Representative images of Alizarin Red staining(left), Oil Red O staining(middle) and β-Gal staining(right) in si*Nsun2*-treated group and the NC group. Scale bar: 100 μm. (R) Quantification of calcification(left), lipid formation(middle) and β-Gal positive cells (right). n=5. Data shown as mean ± SEM. \**P*< 0.05, \*\**P* < 0.01, \*\*\**P* < 0.001 by Student’s t test (B, D, E, I, L, N, R).

It is well established that *Nsun2*-mediated tRNA m⁵C modification in the variable loop region not only affects tRNA stability but also regulates protein translation(44). To confirm global protein translation dynamics, primary BMSCs were transfected with si*Nsun2* or NC interference RNA and then incubated them with O-propargyl-puromycin (OP-puro), which is incorporated into nascent polypeptide chains and can be visualised by fluorescence microscopy. Interestingly, compared to the NC group, we observed a marked reduction in the abundance of nascent polypeptides in the si*Nsun2*-treated group, suggesting a global decrease in protein translation efficiency (Fig. 2M and 2N). Dysregulation of protein translation alters local protein synthesis, which is essential for stem cell function(30, 45). We next evaluated protein synthesis in si*Nsun2*-treated group and NC group through puromycin intake assays. Our investigation revealed that *Nsun2* knockdown led to a reduction in global protein synthesis within si*Nsun2*-treated group. Notably, this reduction was partially reversible through ectopic *Nsun2* overexpression (Fig. 2, O and P). These results further supported the role of tRNA m^5^C in promoting protein synthesis, implying that *Nsun2* might regulate protein translation through tRNA epigenetic modification.

### *Nsun2* depletion shifted lineage fate of BMSCs between osteoblast and adipocyte differentiation

Protein synthesis is a fundamental cellular process essential for all cells. Recent studies have reported that RNA m^5^C methylation modulates translational efficiency and thereby influences stem cell fate determination(30). Transcriptional alterations impacting protein synthesis prompted us to further investigate the role of *Nsun2* in BMSCs differentiation. The primary BMSCs were transfected with si*Nsun2* or NC interference RNA and then cultured in osteogenic and adipogenic induction medium, maintaining standardized incubation conditions to assess lineage-specific differentiation capacity. *Nsun2*-deficient BMSCs exhibited impaired osteogenic differentiation capacity, as shown by reduced mineralization detected by Alizarin Red staining (Fig. 2Q-R), and enhanced adipogenic potential, evidenced by increased lipid accumulation measured by Oil Red O staining (Fig. 2Q-R). Furthermore, *Nsun2* deficiency led to a higher percentage of senescent cells, as indicated by elevated SA-β-galactosidase (β-gal) staining compared to the control group. (Fig. 2Q-R). Our data suggest that *Nsun2* deficiency induces aberrant post-transcriptional m^5^C methylation of tRNA, which may serve as a novel and important contributor to BMSCs senescence and the age-related lineage switch between osteogenic and adipogenic fates.

Next, to identify the mRNA expression profile regulated by *Nsun2*, we isolated total RNA from primary BMSCs that were transfected with si*Nsun2* or NC interference RNA for RNA-seq analysis. A total number of 220 differentially translated mRNAs were identified, including 136 mRNAs with upregulated expression and 84 mRNAs with downregulated expression (Fig. S1A). Gene ontology and pathway analyses showed that the differentially expressed mRNAs were significantly enriched in pathways associated with inflammatory response, regulation of bone remodeling, and aging (Fig. S1B). Notably, loss of *Nsun2* led to a coordinated upregulation of pro-senescent and pro-adipogenic gene programs, including key regulators of cell cycle arrest and lipid metabolism, while genes critical for osteoblast lineage commitment and extracellular matrix mineralization were markedly downregulated (Fig. S1C). And we validated the results by RT-qPCR (Fig. S1D). Together, these findings indicate that *Nsun2* deficiency shifts the osteogenic differentiation of BMSCs toward adipogenic differentiation and promotes cellular senescence through translational regulation.

### *Nsun2*-mediated m⁵C tRNA modification selectively regulates translation of pro-osteogenic factors

Consistent with previous studies, we also support that the lack of *Nsun2* expression broadly impairs protein synthesis(31). Therefore, we conducted LC-MS/MS mass spectrometry to analyze the proteomic profiling of si*Nsun2*-treated group and the NC group. Among a total of 7377 proteins identified in the primary BMSCs, 578 were downregulated and 312 were upregulated in si*Nsun2*-knockout cells compared to the control cells (fold change ≥ 1.2 for upregulation, ≤ 0.83 for downregulation) (Fig. 3A). Subcellular localization analysis revealed that the majority of differentially downregulated proteins (DEPs) were localized to the cytoplasm, suggesting that *Nsun2* depletion predominantly impairs cytoplasmic protein translation (Fig. 3B). Strikingly, GO enrichment analysis revealed that pathways associated with cell differentiation, skeletal system development and bone remodeling were significantly enriched among the list of downregulated proteins (Fig. 3C). The heatmap showing differentially downregulated proteins revealed that a majority of pro-osteogenic factors were significantly decreased in si*Nsun2*-treated group relative to the NC group (Fig. 3D). Compared to osteogenesis differentiation-associated genes identified through RNA-seq analysis, we observed a poor correlation between the proteome and transcriptome, which is consistent with the findings in the *Nsun2*-deficient cortex(37). This discrepancy suggests that the proteomic deficits in *Nsun2*-deficient cells cannot be fully explained by transcriptional changes alone. In light of the results from *Nsun2* deficiency-induced changes in tRNA abundance, we hypothesized that this could alter translational efficiency. We next employed ribosome footprinting (Ribo-Seq) to assess codon usage patterns and relative translation efficiencies (TE) in si*Nsun2*-treated group relative to the NC group (Fig. 3E). Translational efficiency (TE) analysis revealed that 727 mRNAs exhibited a >2-fold increase in TE, while 749 mRNAs showed a <0.5-fold decrease in TE in si*Nsun2* BMSCs (Fig. 3F). Altered tRNA abundance and function is expected to lead to changed ribosome dwell time (DT) at the cognate codon(24). Comparison of codon occupancy revealed that knockdown of *Nsun2* results in increased ribosome dwell time at m^5^C-tRNA-decoded codons, observed at both the A-site and P-site compared to the control BMSCs. This suggests that *Nsun2*-mediated m^5^C tRNA modification is essential for efficient ribosome transition on m^5^C-tRNA-decoded codons. (Fig. 3G). Gene ontology analysis revealed that mRNAs with reduced translation efficiency (TE) in the si*Nsun2*-treated group - functioning in key biological processes including regulation of bone development, bone growth, stem cell proliferation, and stem cell differentiation - were significantly enriched following *Nsun2* depletion. This enrichment pattern underscores *Nsun2*’s pivotal role in maintaining translational control of mRNAs critical for skeletal development and stem cell maintenance (Fig. 3H). Codon pausing combined with proteomics analysis revealed *Nsun2* knockdown significantly elevated codon pausing scores in pro-osteogenic factors (including *Col1a1, Col1a2, Lrp1, Spp1*, *Fn1,* and others) (Fig. 3I). This indicates that these osteogenic factors may experience translational pausing, which could affect their protein expression and functional contributions to bone formation. We next validated protein expression using Western blot analysis. The results demonstrated a significant reduction in protein levels in the si*Nsun2*-treated group relative to the NC group (Fig. 3J and K). Collectively, our findings elucidate the pivotal role of *Nsun2* and the m^5^C tRNA methylome in orchestrating the translational regulation of key pro-osteogenic genes. This discovery highlights a novel epigenetic mechanism governing osteogenic differentiation, where tRNA methylation dynamically modulates the efficiency of ribosomal decoding for specific mRNAs encoding osteogenic transcription factors. The data further establish *Nsun2* as a master regulator of tRNA-mediated translational control in mesenchymal stem cell lineage commitment.

**Figure 3.**
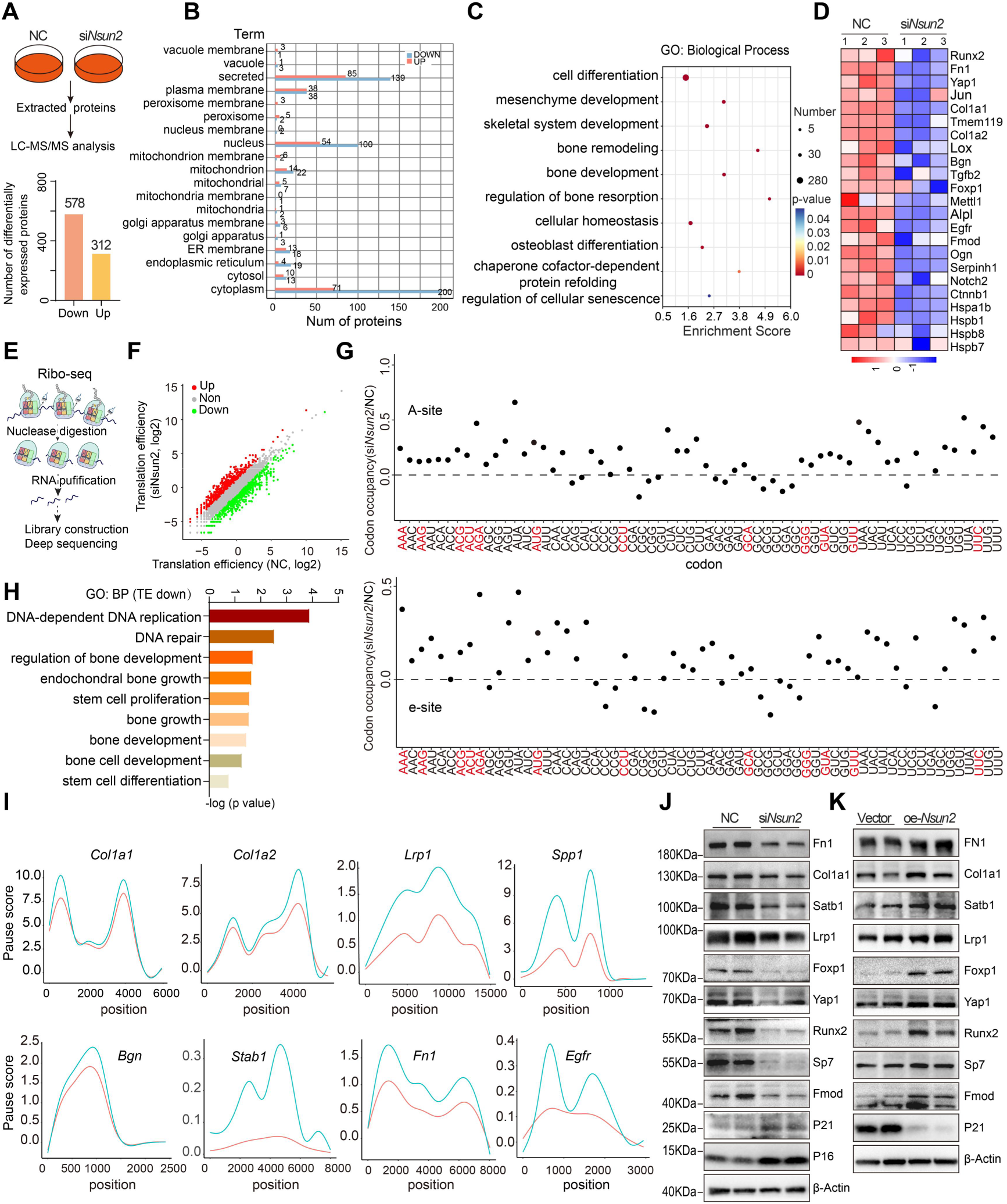
Nsun2-mediated m⁵C tRNA modification selectively regulates translation of pro-osteogenic factors. (A) Schematic diagram of LC-MS/MS mass spectrometry to analyze the proteomic profiling of primary BMSCs transfected with negative control RNA or siRNA of *Nsun2*. (B) Subcellular localization analysis of differentially expressed proteins in si*Nsun2*-treated group relative to the NC group. (C) GO analysis revealed enrichment of biological processes among the differentially expressed proteins. (D) Heatmap of differentially downregulated proteins in si*Nsun2*-treated group relative to the NC group. (E) Schematic diagram of ribosomal footprinting assays (Ribo-seq). (F) Scatterplot of Translation efficiency (TE) from si*Nsun2*-treated group relative to the NC group. (G) Ribosome occupancy at individual codons at A site and E site. Plots represent the relative ribosome protected fragment signals between two groups. (H) Gene ontology analysis revealed enrichment of biological processes among the TE downregulated genes of si*Nsun2*-treated group relative to the NC group. (I) Relative ribosome pausing levels of osteogenic differentiation related genes (*Col1a1*, *Col1a2*, *Lrp1*, *Spp1*, *Bgn*, *Satb1*, *Fn1*, *Egfr*). (J) Representative Western blot images of osteogenic differentiation related proteins and senescence-associated proteins of si*Nsun2*-treated group and the NC group. (K) Representative Western blot images of osteogenic differentiation related proteins and senescence-associated proteins of BMSCs transfected with vector or *Nsun2* plasmid.

### BMSC-specific *Nsun2* deletion impairs bone formation

As efficient protein synthesis specification is indispensable for bone homeostasis, and that translational defects have been increasingly recognized as drivers of impaired osteogenesis and bone loss(46). While our in *vitro* findings strongly implicate *Nsun2* in regulating BMSCs fate decisions, it remained unclear whether such defects are sufficient to impact bone homeostasis in *vivo*. Hence, we generated BMSCs-specific *Nsun2* knockout mice (*Nsun2*^LepR-KO^) by crossing LepR-cre mice with *Nsun2*^flox/flox^ mice. To evaluate the knockout efficiency of *Nsun2* in BMSCs, we isolated BMSCs from young *Nsun2*^LepR-KO^ mice and *Nsun2*^flox/flox^ mice, the knockout efficiency in BMSCs was significant through RT-qPCR assay (Fig. S2A-B). Microcomputed tomography (μCT) analysis of the distal femoral metaphysis showed significantly decreased bone volume, trabecular thickness, and trabecular number, accompanied by a marked increase in trabecular separation in 3-month-old and 13-month-old male *Nsun2*^LepR-KO^ mice compared to *Nsun2*^flox/flox^ littermates (Fig. 4A-E). We next collected femur samples of those mice and stained with LEPR antibody, which represented a reduction in BMSCs populations in the bone marrow from *Nsun2*^LepR-KO^ mice compared to *Nsun2*^flox/flox^ mice (Fig. 4F-G). Meanwhile, the ratio of CD29^+^SCA1^+^ cells in bone marrow of *Nsun2*^LepR-KO^ mice was decreased using flow cytometry analysis (Fig. 4H-I). Besides, we evaluated bone turnover and marrow adiposity. Histological analysis demonstrated a significant reduction in osteoblast number on trabecular surfaces (Fig. 4J-K) and a concomitant increase in adipocyte count within the bone marrow of *Nsun2*^LepR-KO^ mice, compared to their littermate controls (Fig. 4L-N). Calcein double-labeling indicated significantly reduced mineral apposition rate (MAR) and bone formation rate (BFR) in knockout mice (Fig. 4O-P), yet osteoclast numbers were unchanged (Fig. 4Q-R). Likewise, female *Nsun2*^LepR-CKO^ mice displayed significant differences in bone mass, osteoblast number, and increased marrow adiposity compared to littermate controls, while osteoclast number remained unchanged (Fig. S2C-G). Together, these results demonstrate that *Nsun2* deficiency in BMSCs impairs bone formation and accelerates age-associated bone loss.

**Figure 4.**
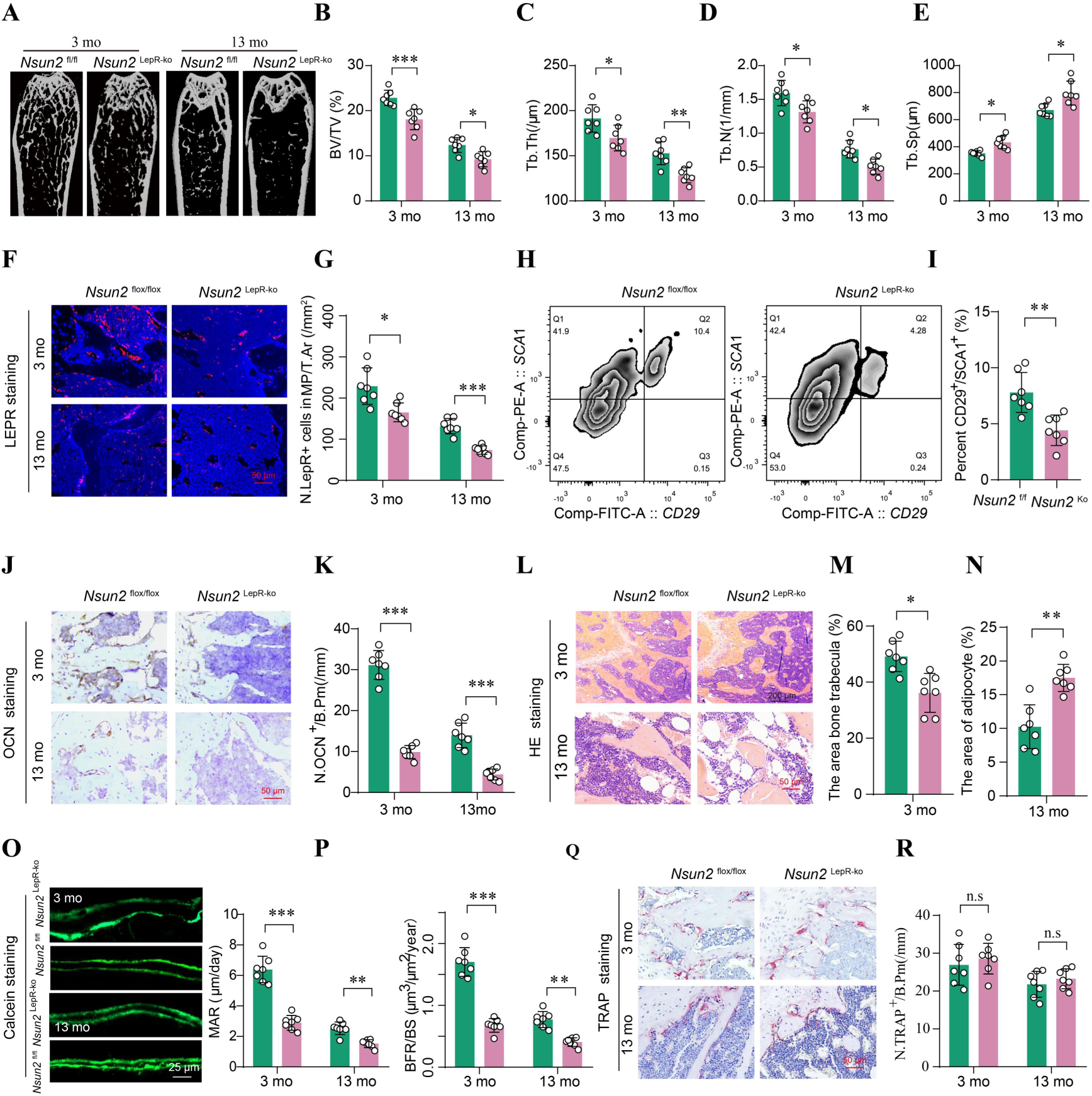
BMSC-specific *Nsun2* deletion impairs bone formation. (A) Representative micro-CT images of trabecular bone in femurs from 3- and 13-month-old male *Nsun2*^flox/flox^ mice and *Nsun2*^LepR-KO^ mice. n=7. (B-E) Quantitative analysis of trabecular bone volume/total volume (BV/TV, B), trabecular thickness (Tb.Th, C), trabecular bone number (Tb.N, D) and trabecular separation (Tb.Sp, E) of femurs. n=7. (F-G) Representative immunofluorescence images of LepR staining (F) and quantification (G) of number of LepR⁺ cells in femurs from 3- and 13-month-old male *Nsun2*^flox/flox^ mice and *Nsun2*^LepR-KO^ mice. Scale bar: 50 μm. n=7. (H-I) Flow cytometry analysis (H) and quantification of CD45^-^CD11b^-^CD29^+^Sca1^+^ cells (I) in bone marrow from 3- and 13-month-old male *Nsun2*^flox/flox^ mice and *Nsun2*^LepR-CKO^ mice. n=7. (J-K) Representative images of osteocalcin staining (J) and quantification (K) of number of osteocalcin^+^ cells in femurs. Scale bar: 50 μm. n=7. (L-N) Representative images of H&E staining (L) and quantification of trabecula area (M) and adipocyte area (N) in femurs, respectively. Scale bar: 200 μm (above), 50 μm (below). n=7. (O and P) Representative images of calcein double labeling(O) and quantification of the mineral apposition rate (MAR, P left panel) and bone formation rate (BFR, P right panel). Scale bar: 25 μm. n=7. (Q and R) Representative images of TRAP staining (Q) and quantification (R) of number of TRAP^+^ cells in femurs. Scale bar: 50 μm. n=7. Data shown as mean ± SEM. n.s > 0.05, \**P* < 0.05, \*\**P* < 0.01, \*\*\**P* < 0.001 by one-way ANOVA (B-E, G, I, K, M-N, P, R).

### Restoration of *Nsun2* enhances BMSCs function and bone formation

According to our results, the m⁵C methyltransferase *Nsun2* exhibited a significant reduction in senescent BMSCs (Fig. 2A-B). *Nsun2* deficiency in these cells correlated with reduced tRNA m⁵C levels, which altered translational efficiency and thereby influenced stem cell differentiation fate while impairing bone formation. Consistent with previous studies, age-associated translational decline and proteostasis collapse are increasingly recognized as key drivers of stem cell dysfunction(2). We therefore hypothesize that restoring *Nsun2* expression could alleviate age-related impairment in bone formation. To this end, BMSCs isolated from aged mice were transfected with *Nsun2*, *Nsun2*-K190M (an enzymatically inactive mutant, in which lysine 190 is substituted by methionine and catalytic activity is abolished)(47), or an empty vector control. Overexpression of *Nsun2* markedly increased mineralized nodule formation, as revealed by Alizarin Red staining, and significantly reduced the proportion of senescence-associated β-gal positive cells (Fig. 5A-D). In parallel, *Nsun2*-transfected BMSCs exhibited fewer protein aggregates together with enhanced nascent peptide synthesis (Fig. 5E-H). These improvements were absent in cells expressing *Nsun2*-K190M, underscoring the requirement for its catalytic activity (Fig. 5A-H). These data demonstrate that *Nsun2* overexpression attenuates cellular senescence and age-related functional deficits by enhancing protein synthesis and reducing protein aggregate accumulation. To extend our in vitro findings, we next assessed the impact of *Nsun2* on BMSCs differentiation and bone formation in vivo. Aged mice (20-month-old) received intramedullary injections of AAV-9 vectors encoding *Nsun2*, *Nsun2*-K190M, or vehicle for six weeks, followed by femoral harvest (Fig. 5I). Although both AAV-*Nsun2* and AAV-*Nsun2*-K190M administration increased *Nsun2* mRNA expression, only *Nsun2* delivery significantly improved bone mass, as evidenced by higher trabecular bone volume, trabecular number, and cortical thickness (Fig. 5J and K). Moreover, *Nsun2*-treated mice exhibited an expansion of LEPR positive BMSCs (Fig. 5L and M) and increased osteoblast numbers on trabecular surfaces without changes in osteoclast abundance (Fig. 5N-Q). Enhanced bone mineralization was further confirmed in the AAV-*Nsun2* group (Fig. 5R-S). In contrast, none of these beneficial effects were observed following AAV-*Nsun2*-K190M treatment (Fig. 5K-S). Overall, these results demonstrate that *Nsun2* promotes osteogenic differentiation and attenuates senescence in BMSCs, thereby enhancing bone formation in aged mice.

**Figure 5.**
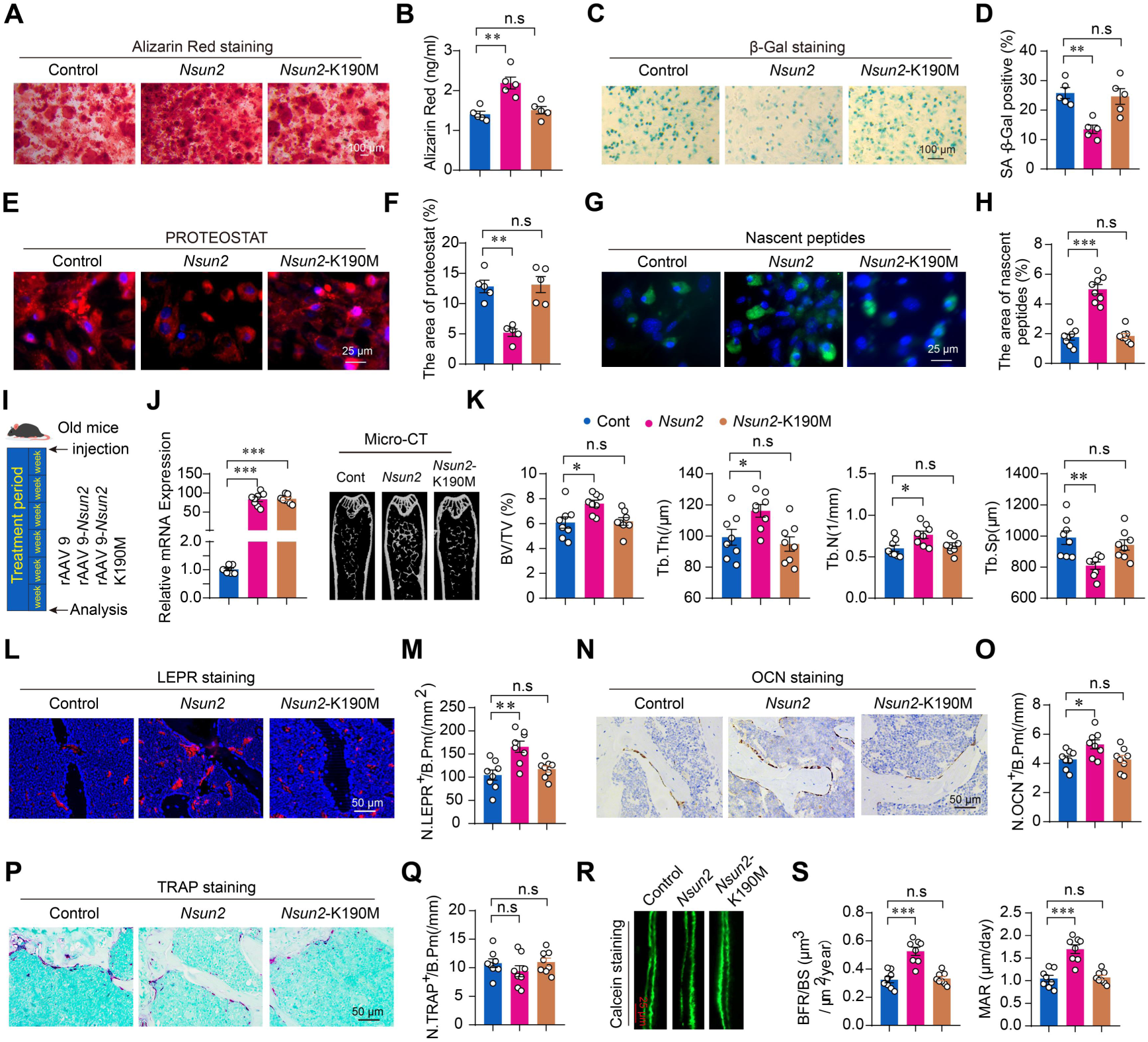
Restoration of *Nsun2* enhances BMSCs function and bone formation. (A and B) Representative images of Alizarin Red staining and quantification of calcification in primary BMSCs transfected with vehicle, *Nsun2* plasmid or *Nsun2*-K190M plasmid. Scale bar: 100 μm. n=5. (C and D) Representative images of β-Gal staining and quantification of β-Gal positive cells of three groups. Scale bar: 100 μm. n=5. (E and F) Representative images of PROTEOSTAT staining and quantification of PROTEOSTAT-positive granules of three groups. Data represent the percentage of PROTEOSTAT-positive granules area. Scale bar: 25 μm. n=5. (G and H) Representative immunofluorescence images and quantification of nascent peptides of three groups. Scale bar: 25 μm. n=8. (I) Schematic illustration showing old mice intrafemoral injected with rAAV 9-*Nsun2*, rAAV 9-*Nsun2* K190M, or vehicle (rAAV 9). (J) Relative mRNA expression of *Nsun2* in old mice following intrafemoral injection of rAAV9 constructs for 6 weeks. n=8. (K) Representative micro-CT images and quantitative of BV/TV, Tb.Th, Tb.N, and Tb.Sp of femurs in old mice intrafemoral injected with rAAV 9-*Nsun2*, rAAV 9-*Nsun2* K190M, or vehicle (rAAV 9). n=8. (L and M) Representative immunofluorescence images of LepR staining (L) and quantification (M) of number of LepR⁺ cells in femurs of old mice. Scale bar: 50 μm. n=8. (N and O) Representative images of osteocalcin staining (N) and quantification (O) of number of osteocalcin^+^ cells in femurs of old mice. Scale bar: 50 μm. n=8. (P and Q) Representative images of TRAP staining (P) and quantification (Q) of number of TRAP^+^ cells in femurs of old mice. Scale bar: 50 μm. n=8. (R and S) Representative images of calcein double labeling(R) and quantification of the MAR(S) and BFR(S) in femurs of old mice. Scale bar: 25 μm. n=8. Data shown as mean ± SEM. n.s > 0.05, \**P* < 0.05, \*\**P* < 0.01, \*\*\**P* < 0.001 by one-way ANOVA (B, D, F, H, J, K, M, O, Q, S).

### *Nsun2* deficiency disrupts proteostasis via impaired molecular chaperone synthesis, triggering ER Stress and UPR activation in BMSCs

Having established that *Nsun2* deficiency impairs translational efficiency and disrupts the protein synthesis of pro-osteogenic factors, which consequently disrupts osteogenic differentiation of BMSCs and induces age-associated bone formation. However, the molecular mechanism by which *Nsun2* deficiency induces BMSC senescence remains unclear. Several studies have reported that tRNA modification defects trigger proteostasis collapse, characterized by pathological protein aggregation, which may be caused by ribosomal pausing and is regarded as a fundamental hallmark of aging and a key driver of age-related diseases(24, 46, 47). Given the hypomodified tRNA in *Nsun2*-deficient cells, we asked whether protein aggregation might also occur in these cells. PROTEOSTAT staining revealed a pronounced increase in global protein aggregates in *Nsun2*-KO cells compared to control cells (Fig. 6A), while *Nsun2*-transfected senescent BMSCs exhibited fewer protein aggregates (Fig. 5E). This is consistent with the findings in senescent BMSCs that senescent cells accumulate protein aggregates (Fig. S3 A-B), indicating that *Nsun2* deficiency induces age-associated protein aggregation, which may serve as a key mechanism driving cellular senescence. The following question is: How does the deletion of *Nsun2* lead to the proteostasis loss. We know that the achievement of proteostasis depends on the proteostasis network (PN) for appropriate protein quality control (PQC), in which the main players are molecular chaperones and two protein degradation systems-the autophagy as well as ubiquitin-proteasome system (UPS)(4). It is well known that aging frequently disrupts the proteostatic equilibrium, shifting it toward pathological states characterized by the accumulation of misfolded protein aggregates, impaired clearance mechanisms, and dysfunctional signaling cascades. This disruption is primarily driven by the dysregulation of chaperones networks and protein degradation machineries(1). Given the positive correlation between cytosine-5 RNA methylation and protein synthesis(31, 32), we hypothesize that *Nsun2* deficiency impairs the synthesis of molecular chaperones, thereby inducing protein aggregation and subsequently disrupting proteostasis in BMSCs. As we expected, our findings indicate that *Nsun2* deficiency reduced the expression of molecular chaperones (including Hspb1, Hspb7, Hspb8, and others) in BMSCs. Ribo-seq confirmed that molecular chaperones *Hspb7* and *Hspb8* exhibited significantly higher codon pausing scores in *Nsun2*-KO cells compared to control cells (Fig. 6B). Concurrently, chaperones including *Hspa4*, *Hspa5*, *Hspa8*, *Hspa9*, and *Hspb8* also exhibited significantly higher codon pausing scores in senescent BMSCs compared to young cells (Fig. S3 C-G). At the protein level, *Nsun2* deficiency suppressed, whereas *Nsun2* overexpression preferentially enhanced, *Hspb7* and *Hspb8* expression (Fig. 6C). These observations supported that *Nsun2* deficiency may impair molecular chaperone synthesis, thereby disrupting proteostasis. As chaperone dysfunction is closely associated with UPR activation and sustained ER stress(48). ER-tracker staining confirmed that *Nsun2* knockout activated ER stress, whereas *Nsun2* overexpression mitigated this effect in senescent BMSCs (Fig. 6D-E). ER stress triggers the activation of the UPR. Western blot demonstrated upregulation of Atf6, Atf4, Ddit3, and p-eIF2α in *Nsun2*-KO cells compared to control cells, indicating that *Nsun2* deficiency directly triggers UPR activation (Fig. 6F). Overall, these results demonstrated that *Nsun2* deficiency impairs molecular chaperone synthesis, leads to misfolded protein aggregation, thereby triggering ER stress and UPR activation, and ultimately causing BMSCs senescence.

**Figure 6.**
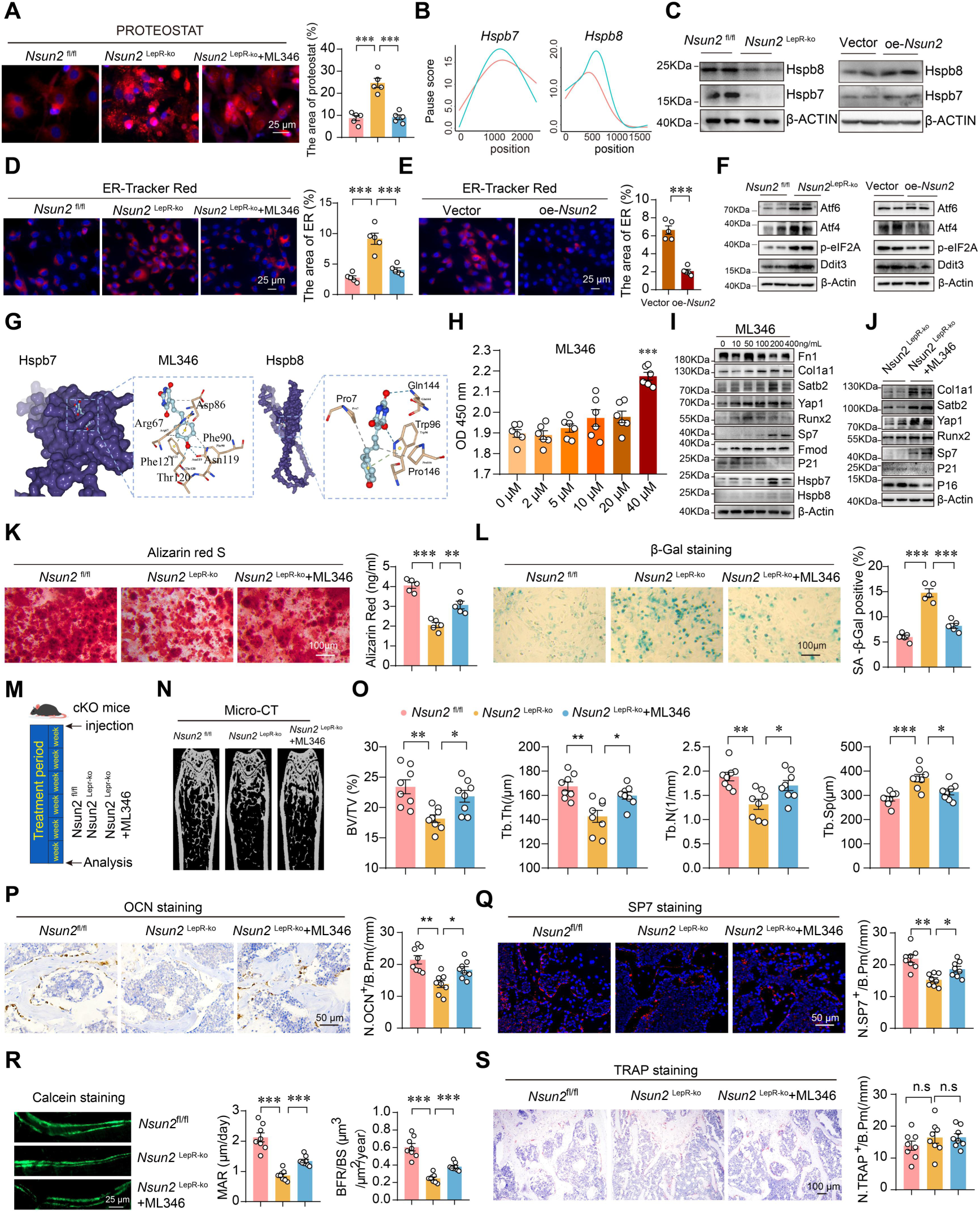
Adminstration of molecular chaperone activator ML346 rescues BMSCs function and bone formation of *Nsun2*^LepR-KO^ mice. (A) Representative images and quantification of PROTEOSTAT (red) and DAPI (blue) staining of BMSCs isolated from *Nsun2*^fl/fl^ or *Nsun2* ^LepR-KO^ mice, followed by the molecular chaperone activator ML346 treatment. Scale bar: 25 μm. n=5. (B) Relative ribosome pausing levels of molecular chaperone genes (*Hspb7*, *Hspb8*) of BMSCs isolated from *Nsun2*^fl/fl^ or *Nsun2* ^LepR-KO^ mice. (C) Representative Western blot images of Hspb8 and Hspb7 of BMSCs isolated from *Nsun2*^fl/fl^ or *Nsun2* ^LepR-KO^ mice (left) and BMSCs transfected with vector or *Nsun2* plasmid (right). (D) Representative immunofluorescence images (left) and quantification (right) of ER-Tracker Red (red) and DAPI (blue) staining of BMSCs isolated from *Nsun2*^fl/fl^ or *Nsun2* ^LepR-KO^ mice, followed by ML346 treatment. Scale bar: 25 μm. n=5. (E) Representative immunofluorescence images (left) and quantification (right) of ER-Tracker Red (red) and DAPI (blue) staining of BMSCs isolated from 24mo mice transfected with vector or *Nsun2* plasmid. Scale bar: 25 μm. n=5. (F) Representative Western blot images of endoplasmic reticulum proteins of BMSCs isolated from *Nsun2*^fl/fl^ or *Nsun2* ^LepR-KO^ mice (left) and BMSCs isolated from 24mo mice transfected with vector or *Nsun2* plasmid (right). n=5. (G) The molecular structure of Hspb7, Hspb8, and a model of the interaction site between Hspb7 or Hspb8 and ML346. (H) Cell proliferation rate assessed using a CCK8 assay after administration of different concentration of ML346. (I) Representative Western blot images of osteogenic differentiation related proteins, senescence-associated proteins and chaperone related proteins of primary BMSCs treated with different concentration of ML346. (J) Representative Western blot images of osteogenic differentiation related proteins and senescence-associated proteins of BMSCs isolated from *Nsun2*^fl/fl^ or *Nsun2* ^LepR-KO^ mice, followed by ML346 treatment (40μM). (K) Representative images of Alizarin Red staining quantification of calcification in BMSCs isolated from *Nsun2*^fl/fl^ or *Nsun2* ^LepR-KO^ mice, and ML346-treated BMSCs isolated from *Nsun2*^LepR-KO^ mice. Scale bar: 100 μm. n=5. (L) Representative images of β-Gal staining and quantification of β-Gal positive cells in BMSCs isolated from *Nsun2*^fl/fl^ or *Nsun2* ^LepR-KO^ mice, and ML346-treated BMSCs isolated from *Nsun2*^LepR-KO^ mice. Scale bar: 100 μm. n=5. (M) Schematic illustration showing *Nsun2*^fl/fl^ or *Nsun2* ^Lepr-KO^ mice, followed by intrafemoral injection of ML346. (N) Representative micro-CT images of trabecular bone in femurs from *Nsun2*^fl/fl^ or *Nsun2* ^LepR-KO^ mice, followed by ML346 treatment. n=8. (O) Quantitative of BV/TV, Tb.Th, Tb.N, and Tb.Sp of femurs. n=8. (P) Representative images of osteocalcin staining and quantification of number of osteocalcin^+^ cells in femurs. Scale bar: 50 μm. n=8. (Q) Representative immunofluorescence images and quantification of number of SP7⁺ cells in femurs. Scale bar: 50 μm. n=8. (R) Representative images of calcein double labeling and quantification of MAR and BFR in femurs. Scale bar: 25 μm. n=8. (S) Representative images of TRAP staining and quantification (Q) of number of TRAP^+^ cells in femurs. Scale bar: 100 μm. n=8. Data shown as mean ± SEM. n.s > 0.05, \**P* < 0.05, \*\**P* < 0.01, \*\*\**P* < 0.001 by one-way ANOVA (A, E, F, H, K, L, O-S).

### *Hspb7/Hspb8* chaperone activation ameliorates protein aggregation and reverses bone loss in *Nsun2*-deficient mice

Studies support that enhanced molecular chaperone expression restores proteostasis across diverse protein misfolding disease models(49), strongly suggesting this approach may serve as a promising therapeutic strategy for osteoporosis. Specifically, ML346 has emerged as a potent small-molecule chaperone agonist, drawing significant interest due to its therapeutic potential in restoring proteostasis. The molecular docking model of ML346 with Hspb7/Hspb8 was generated using AUTODOCK 3.0. According to the model (Fig. 6G), the main-chain oxygen atoms are coordinated by Phe90, Asn119, and Thr120 for Hspb7, while Gln144 and Pro146 stabilize the oxygen atoms for Hspb8. Molecular docking analysis revealed that ML346 binds to both Hspb7 and Hspb8, with a stronger affinity observed for Hspb7. The CCK-8 assay was employed to evaluate the effect of ML346 on cell viability. BMSCs treated with varying concentrations of ML346 exhibited a concentration-dependent promotion of cell proliferation in CCK-8 assays, indicating that ML346 is non-toxic and effectively enhances cell viability (Fig. 6H). To explore whether ML346 is an agonist for Hspb7 and Hspb8 and has a promoting effect on pro-osteogenic factors, we treated BMSCs with varying concentrations of ML346. Western blot analysis revealed that ML346 treatment markedly upregulated the expression of Hspb7, Hspb8, and pro-osteogenic factors, while concomitantly reducing p21 expression, suggesting a dual regulatory role in both osteogenic differentiation and cellular senescence (Fig. 6 I). Meanwhile, ML346 treatment also reversed the deficiency of pro-osteogenic factors and the protein aggregation in *Nsun2*-KO cells, suggesting its potential to restore osteogenic potential in knockout models (Fig. 6 J). To investigate whether ML346 treatment reverses the phenotype of *Nsun2* deficiency, BMSCs derived from wild-type and Nsun2^LepR-ko^ mice were treated with 40μM ML346 and subsequently cultured in osteogenic/adipogenic induction medium. Alizarin Red staining and SA-β-galactosidase staining revealed that ML346 treatment restored both osteogenic differentiation capacity and reversed cellular senescence in *Nsun2*-KO cells (Fig. 6K-L). Encouraged by these in *vitro* results, we next evaluated the role of ML346 in bone formation using an in vivo model. *Nsun2*^LepR-KO^ mice were treated with ML346 (10mg/kg) or vehicle via intraperitoneal injection for six weeks to assess its therapeutic potential (Fig. 6M). Micro-CT analysis demonstrated that ML346 treatment effectively reversed bone loss in *Nsun2*^LepR^-KO mice, as evidenced by significant improvements in trabecular bone parameters: increased bone volume fraction (BV/TV), trabecular number (Tb.N), and trabecular thickness (Tb.Th), coupled with reduced trabecular separation (Tb.Sp) when compared to vehicle-treated controls (Fig. 6 N-O). ML346 administration also enhanced osteogenic differentiation relative to untreated *Nsun2*^LepR-KO^ mice (Fig. 6P-Q). Calcein double-labeling further confirmed a recovery in MAR and BFR in the trabecular bone of the ML346 treatment group (Fig. 6R). In contrast, osteoclast activity remained largely unchanged, as evidenced by comparable numbers of TRAP-positive osteoclasts on bone surfaces across groups (Fig. 6S). These collective findings demonstrate that targeting molecular chaperone modulated ML346 represents a promising therapeutic strategy to counteract bone loss by restoring proteostatic balance and promoting osteogenic function.

### ML346 enhances chaperone activity as a therapeutic strategy to address senile osteoporosis

To further investigate the therapeutic effect of ML346 on age-related osteoporosis, we isolated BMSCs from aged mice and treated them with ML346 or equivalent volume of vehicle. ML346 treatment enhanced osteogenic differentiation and promoted formed protein synthesis, while simultaneously suppressing adipogenic differentiation and reducing cellular senescence (Fig. 7A-D). We then examined the effects of ML346 in *vivo* using aged male and female C57BL/6J mice, which received either ML346 or vehicle control for two months. Micro-CT analysis revealed that ML346-treated mice displayed a notable increase in bone volume, accompanied by greater trabecular thickness, higher trabecular number, and decreased trabecular separation compared with vehicle-treated controls (Fig. 7E-J). Consistently, ML346 administration reduced marrow adipocyte numbers and expanded the pool of multipotent progenitor cells (Fig. 7K-N). Histological evaluation further confirmed a significant rise in osteoblast numbers along trabecular surfaces in ML346-treated mice (Fig. 7O-P). Moreover, calcein double-labeling assays demonstrated enhanced mineral apposition and bone formation rates relative to controls (Fig. 7R-S), whereas osteoclast numbers remained unchanged (Fig. 7T-U). In conclusion, these results demonstrate that ML346 restores BMSCs proteostasis, enhances osteogenic differentiation, and effectively mitigates age-related bone loss.

**Figure 7.**
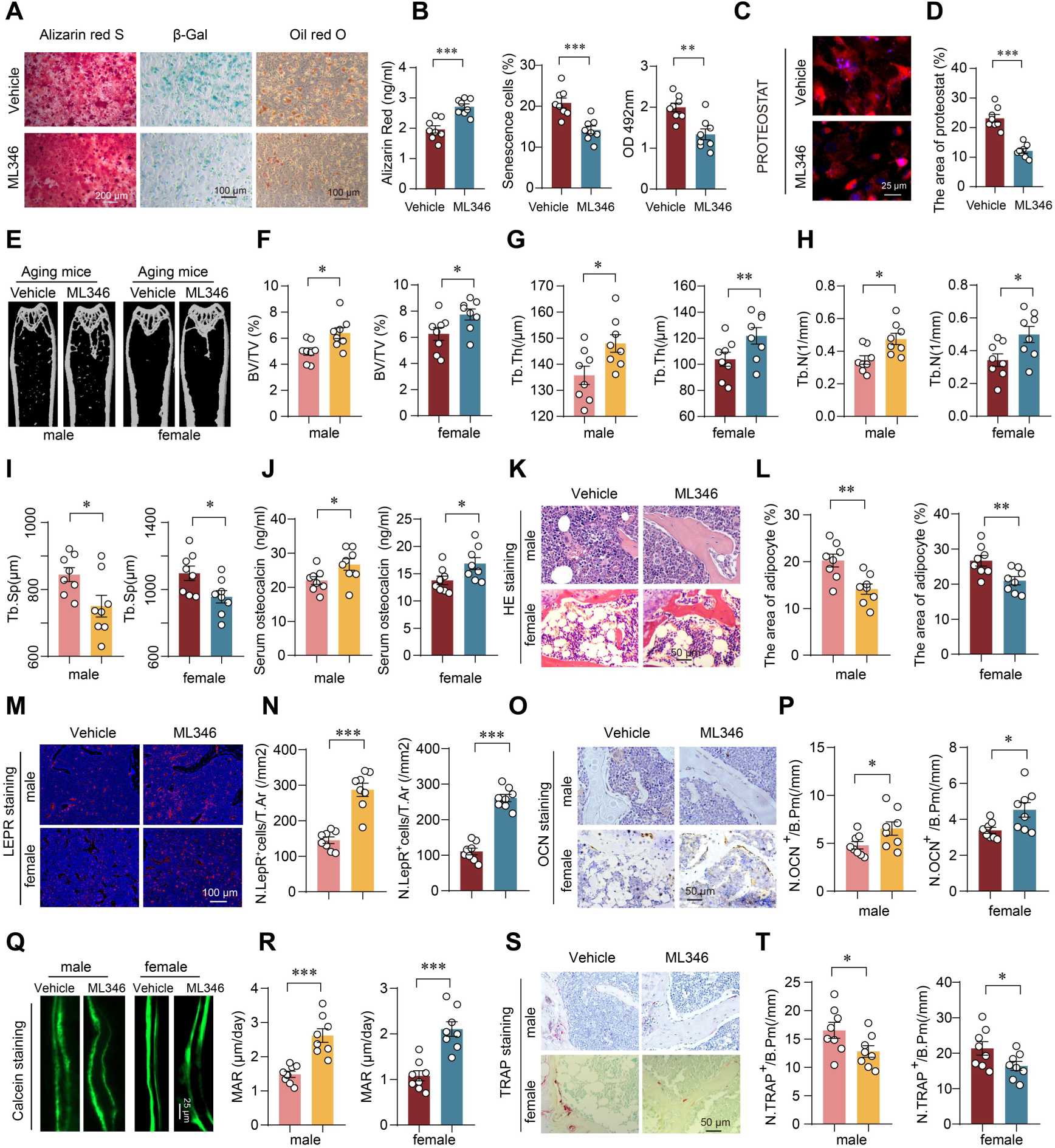
Hspb7/Hspb8 activator ML346 treatment alleviated aging-related bone loss in mice. (A) Representative images of Alizarin Red staining(left), β-Gal staining (middle) and Oil Red O staining (right) in BMSCs isolated from 13-month-old mice treated with ML346 or vehicle. Scale bar: 100 μm. (B) Quantification of calcification(left), β-Gal positive cells (middle) and lipid formation (right) in BMSCs. n=8. (C and D) Representative images (C) and quantification (D) of PROTEOSTAT (red) and DAPI (blue) staining of BMSCs. Scale bar: 25 μm. n=8. (E) Representative micro-CT images of of trabecular bone in femurs from aged male and female mice, followed by ML346 treatment. n=8. (F-J) Quantitative of BV/TV, Tb.Th, Tb.N, and Tb.Sp of femurs. n=8. (K and L) Representative images of H&E staining (K) and quantification of trabecula and adipocyte area (L) in femurs. Scale bar: 50 μm. n=8. (M and N) Representative immunofluorescence images of LEPR staining (M) and quantification (N) of number of LepR⁺ cells in femurs. Scale bar: 100 μm. n=8. (O and P) Representative images of osteocalcin staining(O) and quantification of number of osteocalcin^+^ cells(P) in femurs. Scale bar: 50 μm. n=8. (Q and R) Representative images of calcein double labeling(Q) and quantification(R) of MAR and BFR in femurs. Scale bar: 25 μm. n=8. (S and T) Representative images of TRAP staining(S) and quantification (T) of number of TRAP^+^ cells in femurs. Scale bar: 50 μm. n=8. Data shown as mean ± SEM. \**P* < 0.05, \*\**P* < 0.01, \*\*\**P* < 0.001 by Student’s t test (B, D, F-J, L, N, P, R, T).

## Discussion

*Nsun2* has been primarily studied for its roles in tumorigenesis and neurodegenerative diseases. Here, we reveal its regulatory role in stem cell aging. In aged BMSCs, loss of *Nsun2* destabilizes tRNAs, leading to reduced abundance and impaired chaperone expression, which together compromise proteostasis and aggravate protein aggregation. These findings highlight *Nsun2*-dependent tRNA modifications as a critical mechanism preserving proteostasis and osteogenic potential during aging.

Notably, while previous studies have reported that *Nsun2* promotes osteoclast differentiation by regulating mRNA stabilization and drives bone destruction in tumor-associated bone diseases(50–52). In contrast, our work centers on the impact of *Nsun2* on tRNA stability in aged BMSCs, thereby uncovering a distinct mechanism. To our knowledge, this is the first study investigating tRNA modification in age-related osteoporosis, highlighting the role of *Nsun2* in bone remodeling. However, as prior studies mainly involved tumor-related bone disease. Future work should therefore explore strategies to selectively or transiently enhance NSUN2 activity in bone to mitigate potential adverse effects.

Our findings further expand the current understanding of tRNA modifications in bone. Prior studies on METTL14-, METTL1-, and METTL3-mediated m^5^C modifications primarily emphasize how these modifications orchestrate osteogenic gene expression and cytoskeletal dynamics to promote bone formation or prevent bone loss(53–55) In contrast, our study identifies an additional translational mechanism governed by *Nsun2*-mediated m⁵C modification, which preserves proteostasis by maintaining chaperone synthesis apart from osteogenic protein. Beyond m⁵C, we identified other tRNA modifications, such as m⁶A, which regulates osteoblast and osteoclast differentiation, and m⁷G, which affects bone development, suggesting that multiple tRNA modifications may also contribute to the pathogenesis of osteoporosis(55,56). Interestingly, TRMT6/TRMT61A-mediated m^1^A modification regulates HSC maintenance and self-renewal during aging, whereas our study shows that m⁵C modification can likewise mitigate the age-associated decline of BMSCs(57).This functional conservation suggests that although they are distinct modifications, they mediate highly comparable biological outcomes by preserving stem cell function and counteracting aging. A critical cellular process underpinning stem cell maintenance is energy metabolism, which is governed by mitochondria. As m⁵C modifications are introduced of several mitochondrial tRNAs, where the NSUN2 enzyme partially localizes within mitochondria(58). Consistently, we observed that reduced tRNA modifications were accompanied by a marked decrease in multiple mitochondrial-related proteins, as revealed by proteomic analysis. These findings suggest that regulation of tRNA modification may also influence mitochondrial function. Future studies should therefore explore the interplay between mitochondrial activity and bone metabolism.

During aging, aberrant m^5^C modification in tRNAs can disrupt their stability by reducing binding affinity for m^5^C reader proteins like YBX1(59). This destabilization renders tRNAs more susceptible to cleavage by endonucleases such as angiogenin (ANG)(60), which may represent an important mechanism underlying the age-associated decline in tRNA abundance. Consequently, the process generates tRNA-derived small RNAs (tsRNAs) that actively participate in disease pathogenesis through epigenetic and translational regulation. While their role is well-established in neurodegenerative disorders like PD and AD, the evidence in osteoporosis is still emerging. Nevertheless, dynamic expression of tsRNAs in BMSCs during osteogenic induction and hypoxia implicates them in bone metabolism, influencing differentiation, stress adaptation, and repair(61). Notably, while studies have identified osteoprotective roles for specific tsRNAs, such as tRF-23 promoting osteogenesis in human BMSCs and protecting against bone loss in ovariectomized mice(62), we propose that the production of specific tsRNAs induced by aging may represent a novel mechanism contributing to bone loss. Strikingly, essential amino acids such as lysine not only serve as energy sources but also act as modulators of signaling pathways important for bone homeostasis(63).Based on these observations, supplementation with specific tRNAs or amino acids (e.g Lys, Gly) may help restore age-related osteoporosis.

The effects of tRNA modifications are highly diverse, as they can optimize codon–anticodon interactions, influence translation elongation, and prevent mistranslation(64). Our data further support the significance for tRNA modifications in maintaining translational fidelity, a process known to delay aging and extend lifespan(65). However, it is important to acknowledge that the effects of tRNA modifications are not universally uniform. Certain modifications may slow elongation or induce ribosome pausing, which, rather than compromising fidelity, can selectively reshape the proteome in a context-dependent manner(66). Other tRNA modification enzymes such as METTL1 (m⁷G), TRMT6/61A (m¹A), and the pseudouridine synthases (PUS family) may exert similar fidelity-preserving functions through distinct structural or decoding mechanisms(67–69). We therefore speculate tRNA modification maintains proteostasis by cooperatively regulating translational fidelity, and its decline represents a fundamental molecular basis of aging.

Molecular chaperones play essential roles in maintaining cellular proteostasis by assisting protein folding and facilitating the clearance of misfolded proteins via UPS or autophagy. UPS-mediated ubiquitination governs key signaling pathways in bone cells: it regulates NF-κB signaling and NFATc1 activation in osteoclasts, and fine-tunes osteogenic differentiation in osteoblasts by targeting critical transcription factors for proteasomal degradation(70). These findings are consistent with studies showing that reduced chaperone levels impair osteogenesis and bone homeostasis under pathological conditions(71). In addition to the UPS, autophagy likely contributes, as NSUN2 may regulate protein degradation through interactions with PTEN. Moreover, the HSPB8/BAG3–chaperone-mediated autophagy pathway, which normally promotes the degradation of misfolded proteins, is compromised under NSUN2 deficiency(72). Additional factors, including TRIM family E3 ubiquitin ligases, likely participate in this coordinated degradation network(73). Thus, protein aggregation in bone results not only from increased misfolding but also from failure in clearance mechanisms.

Targeting these pathways offers promising therapeutic opportunities. ML346, belonging to the class of proteostasis regulators (PRs), can restore protein homeostasis by activating HSF-1 and its associated stress response network(49). In addition to ML346, other heat shock protein activators, such as geranylgeranylacetone and celastrol, may similarly restore chaperone activity and protect against age-associated bone loss(74, 75). These findings highlight pharmacological strategies that restore or augment chaperone function as promising therapeutic avenues for osteoporosis. Beyond bone biology, activation of HSPB8 may also mitigate protein aggregation in neurodegenerative diseases such as PD and AD by promoting the clearance of misfolded proteins like α-synuclein and Tau(76), suggesting that ML346 could serve as a potential therapeutic agent for these disorders. Moreover, given the well-established positive correlation between translational fidelity and lifespan extension(77), it is plausible that ML346 may even represent a novel class of longevity-promoting intervention.

Several limitations should be acknowledged in our study. This study mainly focused on the role of tRNA modifications in protein synthesis, while their potential impact on protein degradation pathways, such as the UPS and autophagy, remains unexplored. Future work addressing this gap will help to provide a more integrated view of how tRNA modifications regulate degradation. In summary, NSUN2-dependent tRNA modifications link translational fidelity and proteostasis to bone health, offering insights and potential therapies for age-related osteoporosis.

## Methods

### Animal models

The *Nsun2* conditional knockout mice were generated by Cyagen Company (Suzhou, China). Cas9 protein, two gRNAs (gRNA-B1: TAGATGAATACATACAGGCCAG G, gRNA-B2: ACTTGCTGTCATGTCATGCTGGG) and a donor vector containing two loxP sequences flanked Exon 4∼5 of mouse Nsun2 were co-injected into fertilized eggs. The embryos were transferred to recipient female mice to obtain F0 mice. For genotyping analysis, genomic DNA was isolated from tail tip biopsies using a standardized extraction protocol. PCR amplification was performed with specific genotyping primers for *Nsun2*-floxed mice as follows: forward primer 5’-GAGTCACATCAAAGCCCTTGTTC-3’ and reverse primer 5’-AGCTATGCAGA-CTGAAGAATGAAG-3’. *Nsun2*^flox/flox^ mice were crossed with LepR-Cre mice to generate LepR-Cre^+^; *Nsun2*^flox/flox^ mice, enabling specific knockout of *Nsun2* in bone marrow stromal cells (BMSCs). LepR-Cre mice (catalog C001036) were purchased from Jackson Laboratory (catalog 008320). All mice were maintained on a C57BL/6J genetic background and housed in a specific pathogen-free (SPF) facility at the Laboratory Animal Research Center of Central South University. The housing conditions included a controlled temperature range of 22∼24°C, a 12-hour dark/light cycle, and free access to standard chow and water. Environmental enrichment measures were implemented to promote animal welfare, including cage accessories and nesting materials.

### LC-MS based tRNA modification

Total RNA was extracted from bone marrow stromal cells (BMSCs) derived from 2-month-old and 24-month-old mice with AG RNAex Pro Reagent (AG21102, Accurate Biology), followed by integrity verification via agarose gel electrophoresis and quantification using the Nanodrop™ instrument. Subsequently, tRNA was isolated from total RNA samples by Urea-PAGE electrophoresis and 60∼90nt band of tRNA was excised and purified by ethanol precipitation. The purified tRNA was hydrolysed to single nucleoside and then dephosphorylated in 50 μL reaction volume which containing 10U Benzonase (Sigma), 0.1 U Phosphodiesterase I (US Biological) and 1U Alkaline Phosphatase (NEB) and incubated the reaction at 37 °C for 3 h. Next, pretreated nucleosides solution was deproteinized using Satorius 10,000-Da MWCO spin filter. Analysis of nucleoside mixtures was performed on Agilent 6460 QQQ mass spectrometer with an Agilent 1260 HPLC system using Multi reaction monitoring (MRM) detection mode. LC-MS data was acquired using Agilent Qualitative Analysis software. MRM peaks of each modified nucleoside were extracted and normalized to quantity of tRNA purified.

### tRNA sequencing

Total RNA was extracted from bone marrow stromal cells (BMSCs) derived from three distinct sources - 2-month-old mice, 24-month-old mice, and the si*Nsun2*/Negative Control group - using the AG RNAex Pro Reagent (AG21102, Accurate Biology). From the total RNA, 10 μg was used to isolate small RNAs using the mirVANA miRNA Isolation Kit (Thermo Fisher Scientific; Cat. AM1561). Subsequently, 500 ng of the isolated small RNAs were treated with an AlkB Enzyme Mix (Epibiotek; Cat. R2201) at 37°C for 30 min, after which the RNA was purified using the RNA Clean & Concentrator™-5 Kit (Zymo Research; Cat.R1016). The purified RNA was then sequentially ligated to adapters: first, the 3’ adapter was added and ligated using 3’ Ligase at 28°C for 60 min, followed by a 65°C incubation for 20 min; then, the 5’ adapter was ligated using 5’ Ligase under the same conditions with Epi™ Small RNA Library Fast Kit (Epibiotek; Cat. R2201). Following ligation, an EPI Initiator Mix was added to block excess adapters via gradient cooling. Reverse transcription was then performed by adding a mix containing RT Buffer, RT Enzyme, RT Primer, and RNase Inhibitor, with incubation at 55°C for 60 minutes followed by 70°C for 15 min to synthesize cDNA. The product was immediately purified using Epi™ Small RNA Library Select Beads (Epibiotek; Cat. R2404), and PCR amplification was carried out by adding a 2× HiFi PCR Mix and Index Primers. The final library was purified again using the same beads. Library quality was assessed on a Bioptic Qsep100 Analyzer, and sequencing was performed on a next-generation sequencing (NGS) platform under the help of Aksomics (shanghai). For data Processing, adapter trimming was performed with Cutadapt. The cleaned reads were then aligned to tRNA sequences from the GtRNAdb and mitotRNAdb databases. Differential expression analysis was conducted using DESeq2, with differentially expressed tRNAs defined as those exhibiting an absolute fold change greater than 1.5 and a p-value less than 0.05.

### tRNA bisulfite sequencing

Total RNA was size-selected for small RNA fraction (<200nt) with the MirVana Isolation Kit (ThermoFisher). The enriched small RNAs were demethylated with AlkB enzyme mix for 100 min at 37℃. Demethylated small RNA was bisulfite converted and purified using the EZ RNA methylation Kit (Zymo Research; Cat. R5002). Bisulfite-converted RNA was used for library construction with GenSeq® Small RNA Library Prep Kit (GenSeq, Inc.) by following the manufacturer’s instructions under the help of Cloudseq Biotech Inc. (Shanghai, china). All the libraries were screened for 170∼210bp fragments and qualitied with Qubit3.0. Libraries were sequenced on Illumina NovaSeq with single-end reads. Raw reads were quality controlled by Q30, FastQC, and FastQ Screen. 3’ adaptor- and low quality reads were trimmed by cutadapt software (v1.9.3) to object the high quality clean reads. Meanwhile, the genomic tRNA sequences with high confidence were downloaded from the GtRNAdb database for the corresponding reference genome. A universal CCA triplet was added to the 3’ end of each tRNA sequence in silico to generate tRNA database. Adapter-trimmed and quality-filtered clean reads were then aligned to this tRNA database using meRanGs with default parameters. Subsequent methylation calling was performed using meRanCall to identify m^5^C sites in each sample. For comparative analysis, meRanCompare was employed to identify differentially methylated sites (DMS) between the two experimental groups. A default statistical threshold of a P-value < 0.05 was applied to define significant differential methylation.

### Ribosome sequencing

For ribosome sequencing, BMSCs derived from 2-month-old mice, 24-month-old mice, and the si*Nsun2*/Negative Control group were cultured, followed by the translation blockade using 0.1 mg/mL cycloheximide. Cells were collected and then centrifuged at 500g for 5 min. The cell extracts were resuspended in Polysome Buffer supplemented with 1% Triton X-100, 1 mM DTT, 10 units DNase I, 0.1 mg/mL CHX, and 0.1% NP-40 and incubated on ice for 30 min. After incubation, the lysate was centrifuged at 12,000×g for 10 minutes at 4℃. The supernatant was then collected and mixed with RNase I and DNase I, followed by incubation at room temperature for 45 minutes. 45 min later, the digested reaction was stopped by adding of RNase inhibitor. Subsquently, ribosome protect fragments (RPFs) was isolated using RNA clean & ConcentratorTM-5 kit (Zymo Research; Cat. R1016). EpiTM RiboRNA Depletion Kit (Human/Mouse/Rat) (Epibiotek, R1805) was used for rRNA depletion. Then, RPFs RNA samples were used to construct ribosome sequence libraries with QIAseq miRNA Library kit (QIAGEN; Cat. 1103679) and sequenced on Next-Generation Sequencing (NGS) platform. Adapter sequences were removed from raw sequencing data using cutadapt software. Concurrently, reads of 25∼34 bp in length were aligned to rRNA and tRNA sequences with Bowtie software to exclude ribosomal and transfer RNA reads. The remaining reads were subsequently aligned to the reference genome (Ensembl Version 91) using HISAT2 and to the transcriptome using Bowtie software, respectively. Read counts were calculated using featureCounts software and ORF identification was performed using Price software. The trinucleotide periodicity of ribosomes and codon usage frequency were estimated using revised riboWaltz package. Translation efficiency (TE) was calculated by dividing RPF abundance over the CDS by the mRNA abundance from the input sample. A two-fold change threshold and an FDR < 0.05 were applied to define differentially translated genes. PausePred (https://pausepred.ucc.ie/) was employed to infer ribosome pausing from ribosome profiling data. Peaks in ribosome footprint density were scored based on their magnitude relative to the background density in surrounding regions(78).

### Puromvcin intake assay

Prior to sample collection, the cells were treated with 10 μg/ml puromycin for 20 min and then lysed with RIPA lysis buffer. The samples were separated by SDS-PAGE and electro-transferred to PVDF membranes. Subsequently, the whole membrane was incubated overnight at 4°C with an anti-puromycin antibody (EMD Millipore, 4191133), followed by incubation with secondary antibody the next day. Then the bands were visualized by ChemiDoc XRS Imaging System (Bio-Rad) with chemiluminescence reagent (Thermo Fisher Scientific, 32106). The intensity of the bands corresponds to the amount of puromycin incorporated into the cells: a darker band indicates greater puromycin incorporation, which reflects a higher level of newly synthesized proteins.

### Northern blotting assay

Northern blot was performed using the Biotin Northern Blot Kit (for Small RNA) (R0220, Beyotime) according to the manufacturer’s protocol. Briefly, RNA samples were extracted from BMSCs derived from three distinct sources - 2-month-old mice, 24-month-old mice, and the siNsun2/Negative Control group and then separated on a 12% urea-PAGE gel, followed by transferring the gel to a nylon membrane through semi-dry fast blotter (Guangzhou Dao One, FTB95) at 1.3A for 10 min. Then the membrane was crosslinked under UV irradiation for 5 min. The membrane was pre-hybridized at 37 °C for 2 h, followed by hybridization with a biotin-labeled DNA probe (50 ng/mL) at 37 °C for 2 h. Subsequently, the membrane was fixed 15min and incubated with HRP-streptavidin for 15 min. Signal detection was performed using a chemiluminescent substrate and imaging was visualized by ChemiDoc XRS Imaging System (Bio-Rad). The tRNA probe sequences were listed in Supplemental Table 1.

### PROTEOSTAT® Protein aggregation assay

The primary BMSCs were seeded on glass slides and then washed with PBS and then fixed with 4% formaldehyde for 30 min at room temperatue, followed by permeabilizing with 0.3% Triton X-100 for 5 min. Then, the cells were stained with the PROTEOSTAT dye (1:20,000 dilution, Enzo, ENZ-51023-KP050) for 15 min, washed with 1×PBS 3 times, counterstained with DAPI and fluorescence signals were captured using Zeiss microscope(79).

### Nascent peptides assay

BMSCs were washed and incubated for 30 min with methionine-free medium supplemented with 200 μM L-cystine, 2 mM L-glutamine, 10 mM HEPES, and 50 μM Azide-modified amino acid. Cells were then fixed with 4% PFA for 15 min, permeabilized with 0.5% Triton X-100 for 10 mins, and washed with 1× PBS containing 3% BSA. The Click-iT reaction was performed by incubating cells with a pre-mixed cocktail (100 μL Click-iT® reaction buffer, 10 μL CuSO₄, 10 μL Click-iT^®^ reaction buffer additive 1 solution, and 20 μL Click-iT^®^ reaction buffer additive 2 solution) for 30 min at room temperature, protected from light. Then, cells were incubated with a fluorescently-labeled streptavidin secondary antibody for another 30 min. After final washes, the cells were mounted with mounting medium containing DAPI (10 μg/mL) and imaged.

### Quantitative proteomic profiling based on directDIA

The entire sample was grinded by steel beads with 200 μL of lysis buffer for 20 min, followed by centrifugation at 12,000 × g and 4 °C for 10 min to collect the supernatant. A total of 30 µg of protein from each sample was subjected to reduction and alkylation at 95°C for 5 minutes, digested at 37°C for 2.5 h, purified using a solid-phase extraction cartridge, and finally eluted with peptide elution buffer. The eluted peptides were then vacuum-dried prior to LC-MS/MS analysis. For nanoLC-MS/Msanalysis, peptides (500 ng) were separated using Vanquish neo nano-UPLC system (Thermo Scientific, USA) equipped with an EASY-Spray™ column (150 μm × 15 cm) and a linear gradient of mobile phases A (0.1% formic acid in water) and B (0.1% formic acid in 80% acetonitrile). The eluted peptides were analyzed online using an Astral mass spectrometer (Thermo Scientific, USA) operated in data-independent acquisition (DIA) mode with positive ionization. Full MS scans (m/z 380–980) were acquired at a resolution of 240,000, followed by MS/MS scans (m/z 150–2000) with a 2 m/z isolation window and 25% normalized collision energy. The maximum injection time and AGC target were set to 3 ms and 5×10⁴, respectively. The raw data was analyzed with DIA-NN (v1.8.1) using a species-specific UniProt FASTA database (Mus musculus). Search parameters included: trypsin digestion (max 2 missed cleavages), carbamidomethylation of cysteine as fixed modification, and oxidation of methionine and N-terminal acetylation as variable modifications. Peptide identification was performed with mass tolerances of 20 ppm for both precursor and fragment ions, and results were filtered at 1% FDR at both PSM and peptide levels. Other parameters were kept as default.

### RNA sequencing

Total RNA was isolated from each sample using the RNAiso Plus kit (TAKARA, Japan). RNA quality was examined by gel electrophoresis and with Qubit (Thermo, Waltham, MA, USA). For RNA-sequencing, 1μg of total RNA was used for library construction. Sequencing libraries were generated using VAHTSTM Stranded mRNA-seq Library Prep Kit for Illumina®. The libraries were sequenced as 150bp paired-end reads using Illumina Novaseq X Plus according to the manufacturer’s instructions by the commercial service of Genergy Biotechnology Co. Ltd. (Shanghai, China). The raw data was handled by Skewer and data quality was checked by FastQC v0.11.2 (http://www.bioinformatics.babraham.ac.uk/projects/fastqc/). Clean reads were aligned to the Mouse genome mm10 using STAR. The expression of the transcript was calculated by FPKM (Fragments Per Kilobase of exon model per Million mapped reads) using Perl. Differentially expression analysis between si*Nsun2* and Negative Control group was performed by the R package DESeq2 with the likelihood ratio test option. Differentially expressed genes exhibiting two-fold changes and Benjamini and Hochberg-adjusted P values ≤ 0.05 were selected. Then DEGs were chosen for function and signaling pathway enrichment analysis using GO and KEGG database. The significantly enriched pathways were determined when P <0.05 and at least two affiliated genes were included.

### rAAV9 virus construction and treatment

The *Nsun2* CDS region sequence and the mutated *Nsun2* CDS region (position 569, A→T) were cloned into the pHBAAV-CMV-MSC-3flag-T2A-ZsGreen Shuttle plasmid, or the sh*Hspb8* sequence was cloned into the pHBAAV-U6-MSC-CMV-ZsG-reen vector, and both constructs were transformed into DH5α competent cells for large-scale amplification. These shuttle plasmids were cotransfected into HEK293T cells with pAAV-RC9 and pHelper plasmids. After transfection for 72 hours, HEK293T cells were collected and lysed, and the virus is purified by Adeno-Associated Virus (AAV) Purification Maxi298 Kit (V1469, Biomiga) and then concentrated by ultrafiltration tubes (Millipore, UFC905008). A total of 1*10^11 vg of rAAV9-*Nsun2*, rAAV9-*Nsun2*-K190M, rAAV9-sh*Hspb8* or rAAV9-Null was delivered via periosteal injection into the femoral bone marrow cavity and intervened for 6 weeks.

### BMSCs isolation and culture and transfection

The fresh femurs and tibias isolated from mice in the experimental and control groups were crushed and then digested with collagenase A (Sigma) to obtain a single-cell suspension. The cell suspension was filtered again through a 70μm filter, followed by incubation with FV780 for 20 minutes on ice. Subsequently, the cells were incubated with Sca-1 (BioLegend, 108108, 1:100), CD29 (BioLegend, 102206, 1:100), CD45 (BioLegend, 103132, 1:100), and CD11b (BioLegend, 101226, 1:100) for 20 minutes at 4°C. Cells were then acquired and analyzed on a BD FACS Aria III (BD Biosciences). The Sca-1⁺ CD29⁺ CD45⁻ CD11b⁻ population was sorted as BMSCs. The sorted cells were cultured in α-MEM medium (Gibco) supplemented with 10% fetal bovine serum (FBS, Gibco) and 1% penicillin/streptomycin (Pen/Strep) at 37°C in a humidified incubator with 5% CO₂. For cell function analysis, when BMSCs reached 80%∼85% confluence, each group of cell samples was digested and seeded into culture plates, followed by transfection with siRNA-*Nsun2* (RiboBio Co.) or *Nsun2*-plasmid (Youbio, G145429) using Lipofectamine 2000 (Thermo Fisher Scientific, 11668019) or treatment with ML346 (MedChemExpress, HY-18669). These cells were subjected to induction of adipogenic and osteogenic differentiation.

### Cell differentiation and SA-β-gal staining

For in vitro osteogenic differentiation, BMSCs derived from distinct experimental groups - including 2-month-old mice, 24-month-old mice, *Nsun2*^LepR^-knockout (*Nsun2*^LepR^-ko) mice, rAAV2/8-*Nsun2*-treated mice, rAAV2/8-*Nsun2*-K190M-treated mice, and control mice - as well as BMSCs transfected with sh*Nsun2*, shControl, *Nsun2*-plasmid, *Nsun2*-K190M-plasmid, and empty plasmid vectors - were cultured in α-MEM medium containing 10% fetal bovine serum, 0.1 mM dexamethasone, 10 mM β-glycerophosphate, and 50 μM ascorbic acid-2-phosphate for 21 days. The osteogenic induction medium is replaced every 3 days. After 21 days, the cells were fixed in 4% paraformaldehyde and then stained with 2% Alizarin Red S (OriCell, ALIR-10001) to evaluate extracellular matrix mineralization. A Diaphot Inverted Microscope and Camera System (SunGrant, china) was utilized for imaging. The quantification of Alizarin Red S, released from the extracellular matrix into cetylpyridinium chloride (CPC) solution, was performed spectrophotometrically at 540 nm. For in vitro adipogenic differentiation, the experimental and control BMSCs were cultured in α-MEM containing 10% fetal bovine serum, 0.5 mM 3-isobutyl-1-methylxanthine, 5 μg/ml insulin, and 1 μM dexamethasone for 9 days. Then the Oil Red O staining was performed to detect lipids in mature adipocytes with Oil Red O Solultion (OriCell, OILR-10001). A Diaphot Inverted Microscope and Camera System (SunGrant, china) was utilized for imaging. Oil Red O was extracted from the extracellular matrix and quantified spectrophotometrically at 492 nm. For SA-β-gal staining in vitro, the experimental and control BMSCs were cultured with α-MEM for 3 days. For senescence assessment, cells were fixed and subjected to cellular senescence detection using the Senescence-Associated β-Galactosidase Staining Kit (Solarbio, G1580) following the manufacturer’s instructions. The senescent cells were imaged using the Diaphot Inverted Microscope and Camera System (SunGrant, China). Positive areas were quantified using ImageJ software following standardized protocols.

### Western blot assay

Total proteins were extracted from experimental and control BMSCs using RIPA lysis buffer, separated via SDS-PAGE, and transferred onto PVDF membranes (Millipore). The membranes were then incubated with specific primary and HRP-conjugated secondary antibodies following standardized Western blot protocols. The antibodies used for Western blot are as follows: Hspb8 (Promab, P20444), Hspb7 (Abcam, EPR10106(B)), PUROMYCIN (EMD Millipore, 4191133), Nsun2 (Proteintech, 20854-1-AP), Gapdh (Proteintech, 10494-1-AP), Fn1 (Proteintech, 15613-1-AP), Col1a1 (Proteintech, 14695-1-AP), Satb2 (Promab, P17049), Foxp1 (Abclonal, A9540), Lrp1 (Proteintech, 26106-1-AP), Yap1 (Promab, 30323), Runx2 (Abcam, ab76956), Sp7 (Abcam, AB209484), Fmod (Proteintech, 13281-1-AP), P21 (Abcam, AB109199), P16 (Promab, P21210), Actin (Boster, BM0627), Atf6 (Promab, P22262), Atf4 (Promab, 31162), p-eIF2α (Promab, P20090), Ddi3 (Promab, 30633). Blots were visualized using Basic ECL Electrochemiluminescence Reagent (share-bio, SB-WB001) and imaged with the Chemiluminescence Imaging System (Bio-Rad, USA).

### RT-qRCR assay

Total RNA was extracted from the experimental and control BMSCs using AG RNAex Pro Reagent (AG21102, Accurate Biology), followed by determination of RNA purity and concentration using BioTek Epoch2 microplate reader (BioTek, USA). Then, a total RNA (1μg) was reverse transcribed into cDNA using the 5× Evo M-MLV RT Premix (AG11706, Accurate Biology). Quantitatification of mRNA was detected by ABI QuantStudio 3 system and the relative quantitation value of target genes was given by 2^-ΔΔCT^ method and normalized by GAPDH. Primer sequences are listed in Supplemental Table 2.

### Immunofluorescence and immunohistochemical staining

For immunofluorescence staining, the bones were dissected and fixed with 4% paraformaldehyde for 24h, and then were decalcified in 0.5 M EDTA 21 days at 4 ℃. The bone tissues were then embedded in embedding medium supplemented with 8% gelatin, 2% polyvinylpyrrolidone, and 20% sucrose and sectioned into 6 μm slices, followed by blocking with 3% BSA 1 hour. Then, the sections were incubated with primary antibodies NSUN2 (ThermoFisher Scientific, PA5-112941), Leptin R (AF497, NOVUS) and SP7 (Abcam, AB209484) overnight at 4 ℃. The next day, the samples were treated with Donkey anti-Rabbit IgG (H+L) Alexa Fluor 555 (1:200, Invitrogen, A31572), Donkey anti-Rabbit IgG (H+L) Alexa Fluor 488 (Thermo Fisher Scientific, A21206), Goat anti-Rat IgG (H+L) Alexa Fluor 555 (Thermo Fisher Scientific, A21434) at room temperature for 1h were mounted with glycerol containing DAPI (10 μg/mL), and fluorescence signals were captured using Zeiss microscope (Apotome 3).

For immunohistochemical staining, mouse bones were fixed in paraformaldehyde for 24 h after collection, decalcified at 4°C for 3 weeks, and then embedded in paraffin and sectioned at 6 μm. After deparaffinization and rehydration, the samples were stained with OCN primary antibody (Takara, M042) overnight at 4 ℃, stained by HRP-conjugated secondary antibody 1 h at room temperature, visualized using DAB substrate kit (ZSGB-BIO, PV-8000), and counterstained with hematoxylin.

For tartrate-resistant acid phosphatase (TRAP) staining, which was performed using the Tartrate Resistant Acid Phosphatase (TRAP) Kit (Amizona Scientific, AMK1005-150) according to the manufacturer’s protocol. In brief, after deparaffinization and rehydration, the sections were washed three times with TBS containing 0.5% Tween-20, for 5 min each. Solution 1, Solution 2, Solution 3, and DDH₂O were mixed at a ratio of 4: 4: 28. 5: 368 to prepare the TRAP staining buffer. The mixed solution and the sections were pre-warmed at 37°C for 30 min. Subsequently, the TRAP staining buffer was applied dropwise to cover the bone tissue sections and incubated at 37°C for 15 min. Following incubation, the sections were washed. Solution 4 was then applied to counterstain the nuclei for 15-30 s. After thorough wash, the sections were dehydrated, cleared, and mounted with neutral balsam.

For hematoxylin and eosin (H&E) staining, paraffin-embedded sections were subjected to H&E staining (Servicebio) using the manufacturer’s protocols. Briefly, after deparaffinization and rehydration, the sections were stained with hematoxylin solution for 5 minutes and rinsed with tap water, followed by differentiation in hematoxylin differentiation solution for 2-5 s, and were then blued in hematoxylin bluing solution for 2-5 s with thorough rinsing under running tap water after each step. For eosin staining, the sections were dehydrated through an 85% and 95% graded ethanol series for 5 min respectively. Subsequently, they were stained in eosin solution for 5 min. Finally, after dehydration and clearing, the sections were mounted with neutral balsam. Quantification of male and female mice sections were the percent of bone trabecula area and adipocyte area relative to the tissue area respectively.

### Micro-CT assay

Following dissection, the experimental and control mouse femora were isolated and fixed in 4% paraformaldehyde overnight. The fixed bones were then wrapped in sealing film and scanned on a high-resolution micro-computed tomography (μCT) system (Skyscan 1172, Bruker MicroCT, Kontich, Belgium). Subsequent image reconstruction using NRecon software (v1.6) was followed by analysis in CTAn (v1.9) to quantify trabecular and cortical parameters (trabecular bone volume (BV/TV), trabecular thickness (Tb.Th), trabecular bone number (Tb.N) and trabecular separation (Tb.Sp)), with 3D visualization performed in CTVol (v2.0) for a region extending 5% of the femoral length distal to the growth plate.

### Endoplasmic Reticulum tracker staining

An ER-Tracker Red (beyotime, C1041S) working solution was prepared by diluting the stock 1:1000 in the provided dilution buffer and pre-warmed to 37°C. After treatment, the cell culture medium was removed, and the cells were washed with Hanks’ Balanced Salt Solution (containing Ca²⁺ & Mg²⁺), and then pre-warmed ER-Tracker Red working solution was added and incubated at 37°C for 30 min in the dark. Subsequently, the staining solution was removed, and the cells were washed twice with complete culture medium. Finally, the cells were mounted using glycerin containing DAPI (10 µg/mL) and imaged under a fluorescence microscope.

### Calcein double-labeling assay

To dynamicly assess the bone formation capacity of every batch of experimental mice, we performed a dual calcein labeling protocol,which means that mice were injected with calcein (25 mg/kg, diluted with PBS) on days 8 and 2 before sample collection to create time-marked labels in newly formed bone. After euthanasia, femurs of mice were isolated, fixed in 4% paraformaldehyde overnight, and embedded in methyl methacrylate. Then undecalcified femoral sections with 5 μm thick were cut using a microtome. We evaluated the bone formation ability through two parameters: the mineral apposition rate (MAR), derived by measuring the inter-label distance over the time interval, and the bone formation rate (BFR) which was determined by multiplying MAR by the mineralizing surface per bone surface (MS/BS).

### Statistics

All quantitative data, confirmed to be normally distributed, are expressed as mean ± SEM. These data were analyzed using GraphPad Prism 10. Specifically, two-tailed Student’s t-tests were applied for two-group comparisons, while one-way or two-way ANOVA was employed for multi-group comparisons. Furthermore, all samples were randomly allocated, no data were excluded to maintain objectivity, and all experiments were independently repeated three times with representative results displayed. Statistical significance is consistently denoted across all Fig.s as *p < 0.05, **p < 0.01, and ***p < 0.001.

### Study approval

Animal experiments were performed in compliance with protocols approved by the Subcommittee on Research and Animal Care of Central South University (2024030525).

## Data availability

All the data that support the findings of this study are available from the corresponding author upon reasonable request.

## Author contribution

Y.X. and M.-S.Y. conceived the project and designed the experiments. Z.-L.P. and N.P. performed the experiments. M.-Z.Y. and F.Y. assisted with data acquisition. T.-J.J. and Y.X. performed the data analysis. Y.X. secured funding and provided project administration and supervision. Z.-L.P. and N.P. integrated the data and drafted the manuscript together with Y.X. All authors reviewed and revised the manuscript.

## Acknowledgments

This work was supported by Development Center for Medical Science & Technology National Health Commission of the People’s Republic (2025ZD0550400), National Natural Science Foundation of China (Grant No. 82471619), Natural Science Foundation of Hunan Province (Grant No. 2024JJ5459), and Scientific Research Program of FuRong Laboratory (No. 2024PT5104).

## References

1. Hipp MS, Kasturi P, and Hartl FU. The proteostasis network and its decline in ageing. Nat Rev Mol Cell Biol. 2019;20(7):421–35.

2. Llewellyn J, Hubbard SJ, and Swift J. Translation is an emerging constraint on protein homeostasis in ageing. Trends Cell Biol. 2024;34(8):646–56.

3. Song S, Guo Y, Yang Y, and Fu D. Advances in pathogenesis and therapeutic strategies for osteoporosis. Pharmacol Ther. 2022;237:108168.

4. Ma C, Yu R, Li J, Chao J, and Liu P. Targeting proteostasis network in osteoporosis: Pathological mechanisms and therapeutic implications. Ageing Res Rev. 2023;90:102024.

5. Balchin D, Hayer-Hartl M, and Hartl FU. In vivo aspects of protein folding and quality control. Science. 2016;353(6294):aac4354.

6. Jayaraj GG, Hipp MS, and Hartl FU. Functional Modules of the Proteostasis Network. Cold Spring Harb Perspect Biol. 2020;12(1).

7. Kaushik S, and Cuervo AM. Proteostasis and aging. Nat Med. 2015;21(12):1406–15.

8. Hartl FU, Bracher A, and Hayer-Hartl M. Molecular chaperones in protein folding and proteostasis. Nature. 2011;475(7356):324-32.

9. Kim YE, Hipp MS, Bracher A, Hayer-Hartl M, and Hartl FU. Molecular chaperone functions in protein folding and proteostasis. Annu Rev Biochem. 2013;82:323–55.

10. Sun-Wang JL, Ivanova S, and Zorzano A. The dialogue between the ubiquitin-proteasome system and autophagy: Implications in ageing. Ageing Res Rev. 2020;64:101203.

11. Kaushik S, Tasset I, Arias E, Pampliega O, Wong E, Martinez-Vicente M, et al. Autophagy and the hallmarks of aging. Ageing Res Rev. 2021;72:101468.

12. Wang J, Zhang Y, Cao J, Wang Y, Anwar N, Zhang Z, et al. The role of autophagy in bone metabolism and clinical significance. Autophagy. 2023;19(9):2409–27.

13. Li H, Li D, Ma Z, Qian Z, Kang X, Jin X, et al. Defective autophagy in osteoblasts induces endoplasmic reticulum stress and causes remarkable bone loss. Autophagy. 2018;14(10):1726–41.

14. Huang W, Gong Y, and Yan L. ER Stress, the Unfolded Protein Response and Osteoclastogenesis: A Review. Biomolecules. 2023;13(7).

15. Rattan SI. Synthesis, modification and turnover of proteins during aging. Adv Exp Med Biol. 2010;694:1–13.

16. Tan L, Register TC, and Yammani RR. Age-Related Decline in Expression of Molecular Chaperones Induces Endoplasmic Reticulum Stress and Chondrocyte Apoptosis in Articular Cartilage. Aging Dis. 2020;11(5):1091–102.

17. Alecki C, Rizwan J, Le P, Jacob-Tomas S, Comaduran MF, Verbrugghe M, et al. Localized molecular chaperone synthesis maintains neuronal dendrite proteostasis. Nat Commun. 2024;15(1):10796.

18. Soti C, and Csermely P. Aging and molecular chaperones. Exp Gerontol. 2003;38(10):1037–40.

19. Calderwood SK, Murshid A, and Prince T. The shock of aging: molecular chaperones and the heat shock response in longevity and aging--a mini-review. Gerontology. 2009;55(5):550–8.

20. Ruano D. Proteostasis Dysfunction in Aged Mammalian Cells. The Stressful Role of Inflammation. Front Mol Biosci. 2021;8:658742.

21. Yuan Y, Li J, He Z, Fan X, Mao X, Yang M, et al. tRNA-derived fragments as New Hallmarks of Aging and Age-related Diseases. Aging Dis. 2021;12(5):1304–22.

22. Pereira M, Francisco S, Varanda AS, Santos M, Santos MAS, and Soares AR. Impact of tRNA Modifications and tRNA-Modifying Enzymes on Proteostasis and Human Disease. Int J Mol Sci. 2018;19(12).

23. Liang Y, Ji D, Ying X, Ma R, and Ji W. tsRNA modifications: An emerging layer of biological regulation in disease. J Adv Res. 2025;74:403–14.

24. Nedialkova DD, and Leidel SA. Optimization of Codon Translation Rates via tRNA Modifications Maintains Proteome Integrity. Cell. 2015;161(7):1606–18.

25. Janin M, Coll-SanMartin L, and Esteller M. Disruption of the RNA modifications that target the ribosome translation machinery in human cancer. Mol Cancer. 2020;19(1):70.

26. Torres AG, Batlle E, and Ribas de Pouplana L. Role of tRNA modifications in human diseases. Trends Mol Med. 2014;20(6):306–14.

27. Lu Y, Yang L, Feng Q, Liu Y, Sun X, Liu D, et al. RNA 5-Methylcytosine Modification: Regulatory Molecules, Biological Functions, and Human Diseases. Genomics Proteomics Bioinformatics. 2024;22(5).

28. Song H, Zhang J, Liu B, Xu J, Cai B, Yang H, et al. Biological roles of RNA m(5)C modification and its implications in Cancer immunotherapy. Biomark Res. 2022;10(1):15.

29. Chellamuthu A, and Gray SG. The RNA Methyltransferase NSUN2 and Its Potential Roles in Cancer. Cells. 2020;9(8).

30. Blanco S, Bandiera R, Popis M, Hussain S, Lombard P, Aleksic J, et al. Stem cell function and stress response are controlled by protein synthesis. Nature. 2016;534(7607):335-40.

31. Blanco S, Dietmann S, Flores JV, Hussain S, Kutter C, Humphreys P, et al. Aberrant methylation of tRNAs links cellular stress to neuro-developmental disorders. Embo j. 2014;33(18):2020–39.

32. Gkatza NA, Castro C, Harvey RF, Heiß M, Popis MC, Blanco S, et al. Cytosine-5 RNA methylation links protein synthesis to cell metabolism. PLoS Biol. 2019;17(6):e3000297.

33. Suzuki T. The expanding world of tRNA modifications and their disease relevance. Nat Rev Mol Cell Biol. 2021;22(6):375–92.

34. Dunin-Horkawicz S, Czerwoniec A, Gajda MJ, Feder M, Grosjean H, and Bujnicki JM. MODOMICS: a database of RNA modification pathways. Nucleic Acids Res. 2006;34(Database issue):D145-9.

35. Zhang L, Wei J, Zou Z, and He C. RNA modification systems as therapeutic targets. Nat Rev Drug Discov. 2025.

36. Watanabe K, and Hishiya A. Mouse models of senile osteoporosis. Mol Aspects Med. 2005;26(3):221–31.

37. Blaze J, Navickas A, Phillips HL, Heissel S, Plaza-Jennings A, Miglani S, et al. Neuronal Nsun2 deficiency produces tRNA epitranscriptomic alterations and proteomic shifts impacting synaptic signaling and behavior. Nat Commun. 2021;12(1):4913.

38. Meerson FZ, Javich MP, and Lerman MI. Decrease in the rate of RNA and protein synthesis and degradation in the myocardium under long-term compensatory hyperfunction and on aging. J Mol Cell Cardiol. 1978;10(2):145–59.

39. Cook JR, and Buetow DE. Decreased protein synthesis by polysomes, tRNA and aminoacyl-tRNA synthetases isolated from senescent rat liver. Mech Ageing Dev. 1981;17(1):41–52.

40. Lauria F, Tebaldi T, Bernabò P, Groen EJN, Gillingwater TH, and Viero G. riboWaltz: Optimization of ribosome P-site positioning in ribosome profiling data. PLoS Comput Biol. 2018;14(8):e1006169.

41. Loayza-Puch F, Rooijers K, Buil LC, Zijlstra J, Oude Vrielink JF, Lopes R, et al. Tumour-specific proline vulnerability uncovered by differential ribosome codon reading. Nature. 2016;530(7591):490-4.

42. Frye M, and Blanco S. Post-transcriptional modifications in development and stem cells. Development. 2016;143(21):3871–81.

43. Zhou BO, Yue R, Murphy MM, Peyer JG, and Morrison SJ. Leptin-receptor-expressing mesenchymal stromal cells represent the main source of bone formed by adult bone marrow. Cell Stem Cell. 2014;15(2):154–68.

44. Tuorto F, Liebers R, Musch T, Schaefer M, Hofmann S, Kellner S, et al. RNA cytosine methylation by Dnmt2 and NSun2 promotes tRNA stability and protein synthesis. Nat Struct Mol Biol. 2012;19(9):900–5.

45. Darnell JC. Defects in translational regulation contributing to human cognitive and behavioral disease. Curr Opin Genet Dev. 2011;21(4):465–73.

46. Zhang Y, Kong Y, Zhang W, He J, Zhang Z, Cai Y, et al. METTL3 promotes osteoblast ribosome biogenesis and alleviates periodontitis. Clin Epigenetics. 2024;16(1):18.

47. Hussain S, Benavente SB, Nascimento E, Dragoni I, Kurowski A, Gillich A, et al. The nucleolar RNA methyltransferase Misu (NSun2) is required for mitotic spindle stability. J Cell Biol. 2009;186(1):27–40.

48. Sepulveda D, Rojas-Rivera D, Rodríguez DA, Groenendyk J, Köhler A, Lebeaupin C, et al. Interactome Screening Identifies the ER Luminal Chaperone Hsp47 as a Regulator of the Unfolded Protein Response Transducer IRE1α. Mol Cell. 2018;69(2):238–52.e7.

49. Calamini B, Silva MC, Madoux F, Hutt DM, Khanna S, Chalfant MA, et al. Small-molecule proteostasis regulators for protein conformational diseases. Nat Chem Biol. 2011;8(2):185–96.

50. Zhang M, Tang C, Li S, Jiang X, Li B, Chen Y, et al. NSUN2-mediated m(5)C modification of KDM6B mRNA enhances osteoclast differentiation and promotes breast cancer bone metastasis. Cancer Lett. 2025;631:217939.

51. Yang M, Wei R, Zhang S, Hu S, Liang X, Yang Z, et al. NSUN2 promotes osteosarcoma progression by enhancing the stability of FABP5 mRNA via m(5)C methylation. Cell Death Dis. 2023;14(2):125.

52. Yu M, Cai Z, Zhang J, Zhang Y, Fu J, and Cui X. Aberrant NSUN2-mediated m5C modification of exosomal LncRNA MALAT1 induced RANKL-mediated bone destruction in multiple myeloma. Commun Biol. 2024;7(1):1249.

53. He M, Lei H, He X, Liu Y, Wang A, Ren Z, et al. METTL14 Regulates Osteogenesis of Bone Marrow Mesenchymal Stem Cells via Inducing Autophagy Through m6A/IGF2BPs/Beclin-1 Signal Axis. Stem Cells Transl Med. 2022;11(9):987–1001.

54. Huang M, Xu S, Liu L, Zhang M, Guo J, Yuan Y, et al. m6A Methylation Regulates Osteoblastic Differentiation and Bone Remodeling. Front Cell Dev Biol. 2021;9:783322.

55. Li Q, Jiang S, Lei K, Han H, Chen Y, Lin W, et al. Metabolic rewiring during bone development underlies tRNA m7G-associated primordial dwarfism. J Clin Invest. 2024;134(20).

56. Li D, Cai L, Meng R, Feng Z, and Xu Q. METTL3 Modulates Osteoclast Differentiation and Function by Controlling RNA Stability and Nuclear Export. Int J Mol Sci. 2020;21(5).

57. Zuo H, Wu A, Wang M, Hong L, and Wang H. tRNA m(1)A modification regulate HSC maintenance and self-renewal via mTORC1 signaling. Nat Commun. 2024;15(1):5706.

58. Shinoda S, Kitagawa S, Nakagawa S, Wei FY, Tomizawa K, Araki K, et al. Mammalian NSUN2 introduces 5-methylcytidines into mitochondrial tRNAs. Nucleic Acids Res. 2019;47(16):8734–45.

59. Zhang J, Che P, Yang Z, Zhang P, Shui Y, Lu X, et al. The m5C reader Ybx1 regulates embryonic cortical neurogenesis by promoting progenitor cell cycle progression. PLoS Biol. 2025;23(5):e3003175.

60. Li D, Gao X, Ma X, Wang M, Cheng C, Xue T, et al. Aging-induced tRNA(Glu)-derived fragment impairs glutamate biosynthesis by targeting mitochondrial translation-dependent cristae organization. Cell Metab. 2024;36(5):1059–75.e9.

61. Zhang C, Ye W, Zhao M, Xia D, and Fan Z. tRNA-derived small RNA changes in bone marrow stem cells under hypoxia and osteogenic conduction. J Oral Rehabil. 2023;50(12):1487–97.

62. Liao H, Li W, Xu L, Zhao C, Li X, Xiong J, et al. The novel tRF-23 promotes osteogenic differentiation of hBMSCs and protects against bone loss in ovariectomized mice. Stem Cell Reports. 2025;20(10):102673.

63. Lv Z, Shi W, and Zhang Q. Role of Essential Amino Acids in Age-Induced Bone Loss. Int J Mol Sci. 2022;23(19).

64. Agris PF, Narendran A, Sarachan K, Väre VYP, and Eruysal E. The Importance of Being Modified: The Role of RNA Modifications in Translational Fidelity. Enzymes. 2017;41:1–50.

65. Martinez-Miguel VE, Lujan C, Espie-Caullet T, Martinez-Martinez D, Moore S, Backes C, et al. Increased fidelity of protein synthesis extends lifespan. Cell Metab. 2021;33(11):2288–300.e12.

66. Manickam N, Joshi K, Bhatt MJ, and Farabaugh PJ. Effects of tRNA modification on translational accuracy depend on intrinsic codon-anticodon strength. Nucleic Acids Res. 2016;44(4):1871–81.

67. Lin S, Liu Q, Lelyveld VS, Choe J, Szostak JW, and Gregory RI. Mettl1/Wdr4-Mediated m(7)G tRNA Methylome Is Required for Normal mRNA Translation and Embryonic Stem Cell Self-Renewal and Differentiation. Mol Cell. 2018;71(2):244–55.e5.

68. Wang Y, Wang J, Li X, Xiong X, Wang J, Zhou Z, et al. N(1)-methyladenosine methylation in tRNA drives liver tumourigenesis by regulating cholesterol metabolism. Nat Commun. 2021;12(1):6314.

69. Jürgenstein K, Tagel M, Ilves H, Leppik M, Kivisaar M, and Remme J. Variance in translational fidelity of different bacterial species is affected by pseudouridines in the tRNA anticodon stem-loop. RNA Biol. 2022;19(1):1050–8.

70. Hino S, Kondo S, Yoshinaga K, Saito A, Murakami T, Kanemoto S, et al. Regulation of ER molecular chaperone prevents bone loss in a murine model for osteoporosis. J Bone Miner Metab. 2010;28(2):131–8.

71. Wu T, Jiang Y, Shi W, Wang Y, and Li T. Endoplasmic reticulum stress: a novel targeted approach to repair bone defects by regulating osteogenesis and angiogenesis. J Transl Med. 2023;21(1):480.

72. Carra S, Seguin SJ, and Landry J. HspB8 and Bag3: a new chaperone complex targeting misfolded proteins to macroautophagy. Autophagy. 2008;4(2):237–9.

73. Zhang L, Afolabi LO, Wan X, Li Y, and Chen L. Emerging Roles of Tripartite Motif-Containing Family Proteins (TRIMs) in Eliminating Misfolded Proteins. Front Cell Dev Biol. 2020;8:802.

74. Westerheide SD, Bosman JD, Mbadugha BN, Kawahara TL, Matsumoto G, Kim S, et al. Celastrols as inducers of the heat shock response and cytoprotection. J Biol Chem. 2004;279(53):56053–60.

75. Endo S, Hiramatsu N, Hayakawa K, Okamura M, Kasai A, Tagawa Y, et al. Geranylgeranylacetone, an inducer of the 70-kDa heat shock protein (HSP70), elicits unfolded protein response and coordinates cellular fate independently of HSP70. Mol Pharmacol. 2007;72(5):1337–48.

76. Bonavita R, Vitale F, Verdicchio LV, Williams SV, Caporaso MG, Fleming A, et al. Small HSPs at the crossroad between protein aggregation, autophagy and unconventional secretion: clinical implications and potential therapeutic opportunities in the context of neurodegenerative diseases. Front Cell Dev Biol. 2025;13:1538377.

77. Zheng B, Zhang W, Yu G, Shi W, Deng S, Zhang X, et al. Translational fidelity and longevity are genetically linked. Nat Commun. 2025;16(1):7521.

78. Orellana EA, Liu Q, Yankova E, Pirouz M, De Braekeleer E, Zhang W, et al. METTL1-mediated m(7)G modification of Arg-TCT tRNA drives oncogenic transformation. Mol Cell. 2021;81(16):3323–38.e14.

79. Lv X, Lu X, Cao J, Luo Q, Ding Y, Peng F, et al. Modulation of the proteostasis network promotes tumor resistance to oncogenic KRAS inhibitors. Science. 2023;381(6662):eabn4180.

