## Supplemental Figures for "Aberrant 5-Methylcytosine tRNA Modification Disrupts Proteostasis and Exacerbates Age-Related Osteoporosis"

**Supplementary Figure 1**

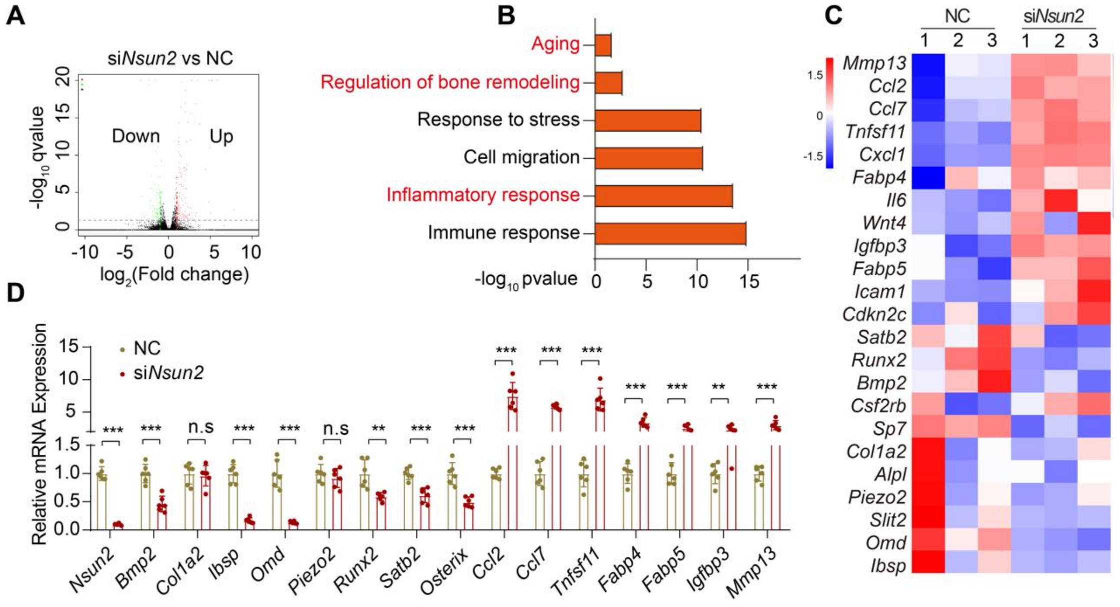


**Supplemental Figure 1.**

(A) Volcano plot of differentially expressed genes in BMSCs transfected with negative control RNA or siRNA of *Nsun2* by RNA-seq.

(B) GO enrichment analysis of differentially expressed genes.

(C) Heatmap of differentially expressed genes involved in the regulation of senescence, inflammation, adipogenic and osteogenic differentiation.

(D) RT-qPCR analysis of genes regulating senescence, inflammation, adipogenic and osteogenic differentiation. n=6.

Data shown as mean ± SEM. ns > 0.05, ***P* < 0.01, ****P* < 0.001 by Student's t test (D)

**Supplementary Figure 2**

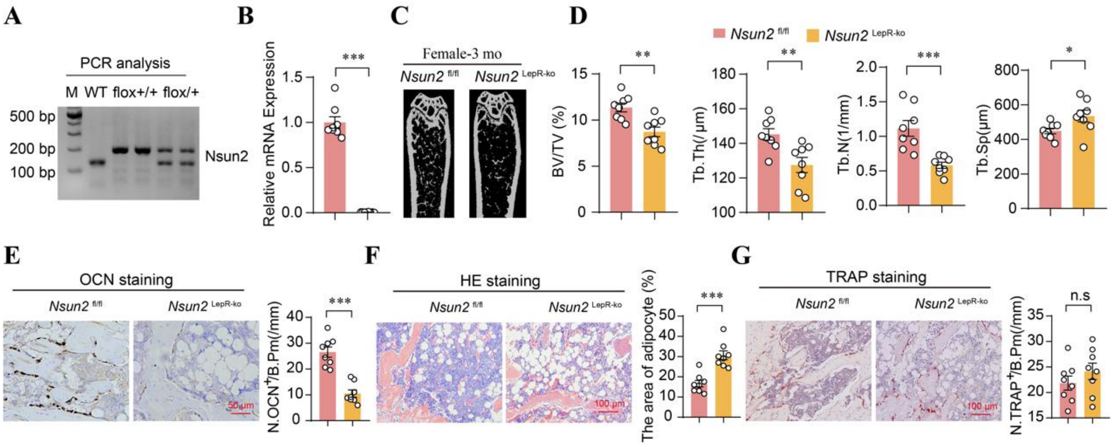


**Supplemental Figure 2.**

(A) PCR analysis of *Nsun2* of wildtype mice, *Nsun2*$^{flox/flox}$ mice and *Nsun2*$^{LepR\text{-}KO}$ mice.

(B) RT-qPCR analysis of *Nsun2* in BMSCs isolated from 3-month-old female *Nsun2*$^{flox/flox}$ mice and *Nsun2*$^{LepR\text{-}KO}$ mice.

(C) Representative micro-CT images of trabecular bone in femurs from 3-month-old female *Nsun2*$^{flox/flox}$ mice and *Nsun2*$^{LepR\text{-}KO}$ mice. n=8.

(D) Quantitative analysis of trabecular bone volume (BV/TV), trabecular thickness (Tb.Th), trabecular bone number (Tb.N) and trabecular separation (Tb.Sp) of femurs. n=8.

(E) Representative images of osteocalcin staining and quantification of number of osteocalcin$^{+}$ cells in femurs. Scale bar: 50 μm. n=8.

(F) Representative images of H&E staining and quantification of adipocyte area in femurs. Scale bar: 100 μm, n=8.

(G) Representative images of TRAP staining and quantification of number of TRAP$^{+}$ cells in femurs. Scale bar: 100 μm. n=8.

Data shown as mean ± SEM. ns ＞ 0.05, *$P$ < 0.05, **$P$ < 0.01, ***$P$ < 0.001 by Student's t test (B, D-G).

**Supplementary Figure 3**

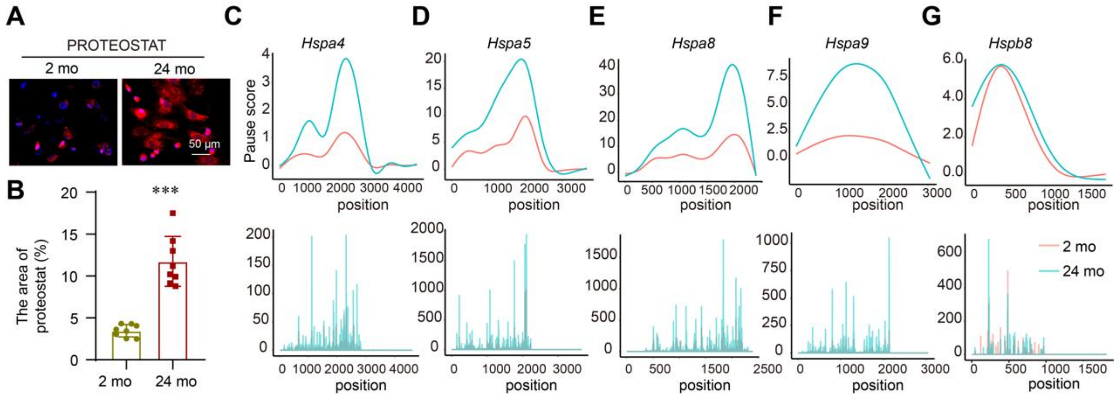


**Supplemental Figure 3.**

(A and B) Representative images and quantification of PROTEOSTAT (red) and DAPI (blue) staining of BMSCs isolated from 2- and 24-month-old mice. Scale bar: 50 μm, n=8.

(C)Relative ribosome pausing levels of *Hspa4* of BMSCs isolated from 2- and 24-month-old mice.

(D) Relative ribosome pausing score of *Hspa5*.

(E) Relative ribosome pausing score of *Hspa8*.

(F) Relative ribosome pausing score of *Hspa9*.

(G) Relative ribosome pausing score of *Hspb8*.
